# Five years of high-frequency data of phytoplankton zooplankton and limnology from a temperate eutrophic lake

**DOI:** 10.1101/2025.03.20.644357

**Authors:** Stefanie Eyring, Marta Reyes, Ewa Merz, Marco Baity-Jesi, Pinelopi Ntesika, Christian Ebi, Stuart Dennis, Francesco Pomati

**Author notes:** Corresponding authors /.

## Abstract

This study presents a comprehensive dataset from Lake Greifen, Switzerland, collected between April 2018 and June 2023, using high-frequency automated monitoring systems. The dataset integrates meteorological data, nutrient chemistry, water column profiles for water physics, and plankton underwater imaging, offering insights into the lake’s physical and biological processes. A dual-magnification dark field underwater microscope captured hourly plankton dynamics at 3 m depth, providing size, shape, and taxonomic information. A profiler with a multiparametric probe monitored water temperature, oxygen, and other key parameters from 1 to 17 m depth, while weekly nutrient sampling complemented the measurements. Data processing involved rigorous cleaning protocols to remove technical artefacts, ensuring data quality. Our dataset showcases the utility of integrating different approaches for high-frequency monitoring to detect lake temporal processes, from phytoplankton blooms to zooplankton vertical migration and seasonal shifts in water column stability. This dataset provides a unique resource for studying limnology and plankton community ecology. All data and related processing codes are publicly available for further research, supporting interdisciplinary studies.

## Background & Summary

Lakes and freshwater reservoirs are critical biodiversity hotspots and provide vital ecosystem services, including drinking water, fisheries, and economic and cultural support. They are also sensitive indicators of human impact ^1^. Climate change, which stresses mountain snow reservoirs ^2^ and groundwater resources ^3^, further enhances the dependence of human society and the local economy on these fragile ecosystems. Understanding and predicting the effects of global change on lakes and reservoirs largely relies on monitoring plankton biodiversity and water physicochemical parameters. Due to their short generation times and rapid response to environmental changes, plankton abundance and diversity are commonly used to assess ecosystem health and water quality ^4–7^.

Significant gaps remain in our understanding of the relative importance of biotic and abiotic controls in the dynamics of lakes and reservoirs, particularly at the scale that matters to predict rapid changes in water quality like algal blooms, which occur within days and weeks due to the fast generation time of plankton, and the rapid hydrodynamic responses to changes in meteorological conditions ^8^ This gap has significant implications for managing water quality and ecosystem services, which requires rapid decision-making. Additionally, understanding ecological processes is crucial for predicting the impact of anthropogenic stressors on biodiversity in freshwater ecosystems ^9,10^.

Traditionally, microscopic identification and analysis have been the primary methods for studying plankton, though they require significant time and expertise from trained taxonomists. The quality of these analyses varies due to human error, which is not easily traceable ^11–14^. The frequency provided by traditional sampling, microscopy and laboratory analysis is insufficient to capture and model lake biogeochemical processes ^15^. Most long-term lake monitoring programs operate at a fortnightly or monthly sampling interval ^15,16^. Particularly for plankton, recent research emphasises the need to quantify both division and loss processes or their net effects at a daily scale when investigating environmental drivers of phytoplankton dynamics ^17–19^. To fill this methodological gap, *in-situ* plankton monitoring technologies like the underwater Dual-magnification Scripps Plankton Camera (DSPC) imaging system offer high-frequency automated plankton monitoring in freshwater ecosystems ^11^. One such DSPC was installed in Lake Greifen (Greifensee), Switzerland in 2018, on a monitoring platform equipped with a meteorological station and an automated water profiler for water physics and chemistry ^15^.

Lake Greifen, a monomictic peri-alpine lake covering 8.45 km^2^, underwent significant eutrophication and subsequent reoligotrophication. During the 1960s and 1970s, the lake’s phosphorus load peaked at 0.6 mg P/L ^20,21^. By 2019, the annual average phosphorus load had decreased to just below 0.05 mg P/L, which still classifies it as eutrophic according to OECD guidelines ^22^. Along with nutrient reductions, rising temperatures have impacted the lake’s limnology (it used to be a dimictic lake) and plankton community. Over the past 50 years, Lake Greifen’s physical, chemical, and planktonic properties have been monitored monthly ^16,20^.

This article presents a five-year high-frequency time series from the monitoring platform located at the northern end of Lake Greifen, featuring fully automated limnology ^11,15^. The station is 450 metres from the shore and 1 km from the lake’s outlet (coordinates: 47° 21’ 59” N, 8° 39’ 54.4” E) at 435 metres above sea level. The depth at the monitoring site is 20 metres, with a maximum lake depth of 32 metres (see SI **Tab. S1** and **Fig. S1** for lake bathymetry). The platform is equipped with a meteorological station, a CTD probe and profiler, and the Dual-magnification Scripps Plankton Camera (DSPC). Additionally, we conduct sampling for nutrient chemistry at a (bi-)weekly scale (**Fig. 1**).

**Fig. 1.**
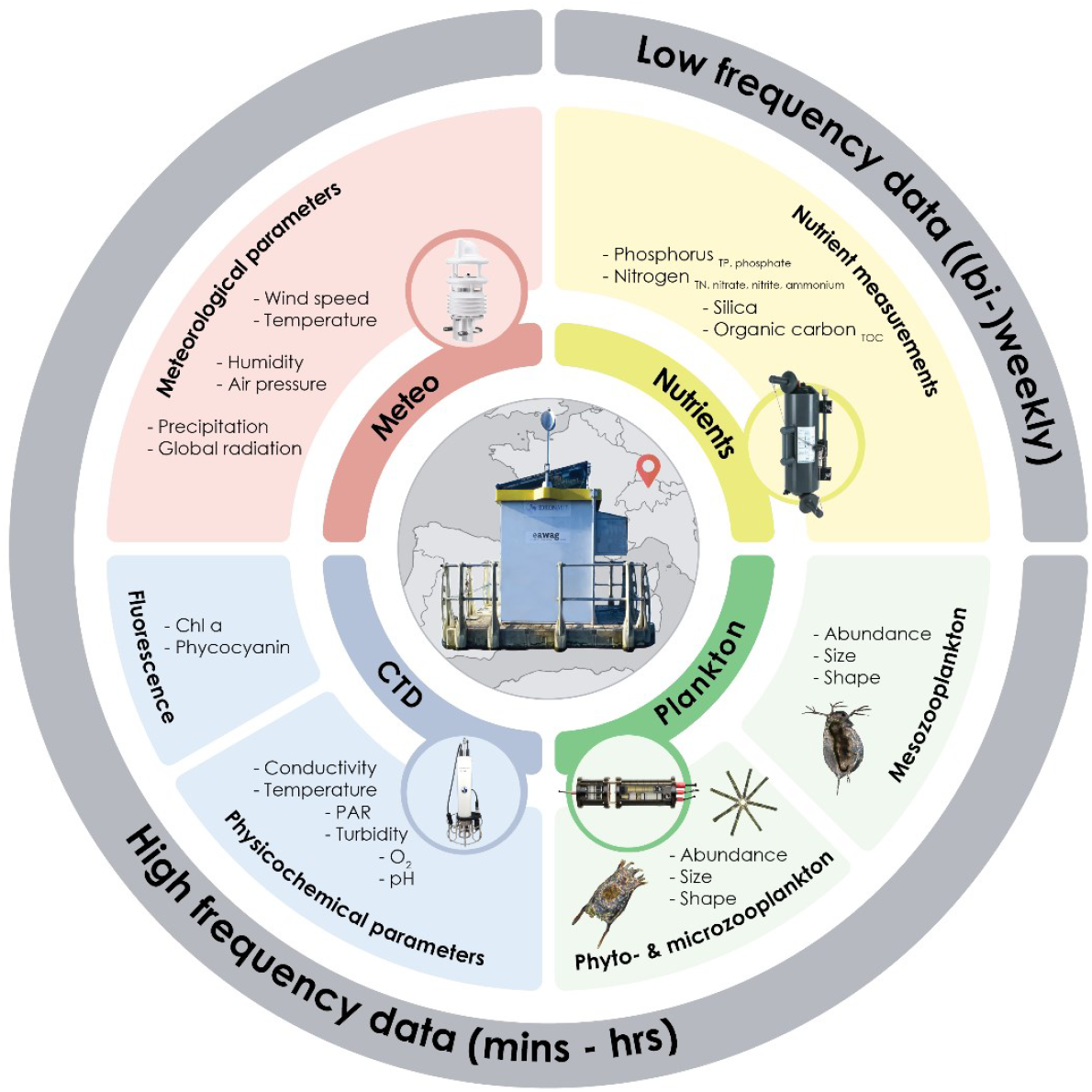
The data: Overview of water analyses conducted, and parameters observed. Figure inspired by ^37^.

## Methods

### Data collection

#### Monitoring platform

The Lake Greifen monitoring platform is powered by solar panels and a fuel cell and connected to a server at Eawag through a cellular router. It is equipped with a computer and a controller system as described in ^15^. A CTD probe, a meteorological station and a DSPC are installed at the platform for automated high-frequency monitoring. In the following, we will discuss each instrument. For an overview of measurement intervals, timezones and time ranges of all instruments installed at the monitoring platform in Lake Greifen consult SI **Tab. S2**.

#### Meteorological station

The instrument with the highest measurement frequency and a measurement interval of 1 to 30 minutes is the meteorological station. We collected data from 18 April 2018 until 30 June 2023 with a larger gap in the first year from 5 December 2018 to 10 May 2019 which is attributed to a exchange of the meteorological station. The meteorological station on the measuring platform was a Vaisala Oyj WXT520 until 2018, which was replaced by a LUFFT (OTT HydroMet brand) WS700-UMB in 2019. The WS700-UMB was initially operated with a Meteobridge PRO data logger until December 11, 2020, which was then replaced by an OTT HydroMet netDL1000. The resolution of the sensors can be found in the manufacturer’s manual. The meteorological station is installed on the roof of the platform at 3.15 m height above the water level. The parameters measured are precipitation (cumulative absolute rainfall, precipitation intensity and precipitation type), global radiation, wind speed (average and maximum wind speed as well as wind direction), air temperature, humidity (relative and absolute), dew point and atmospheric pressure (absolute and relative) (**Fig. 2**).

**Fig. 2:**
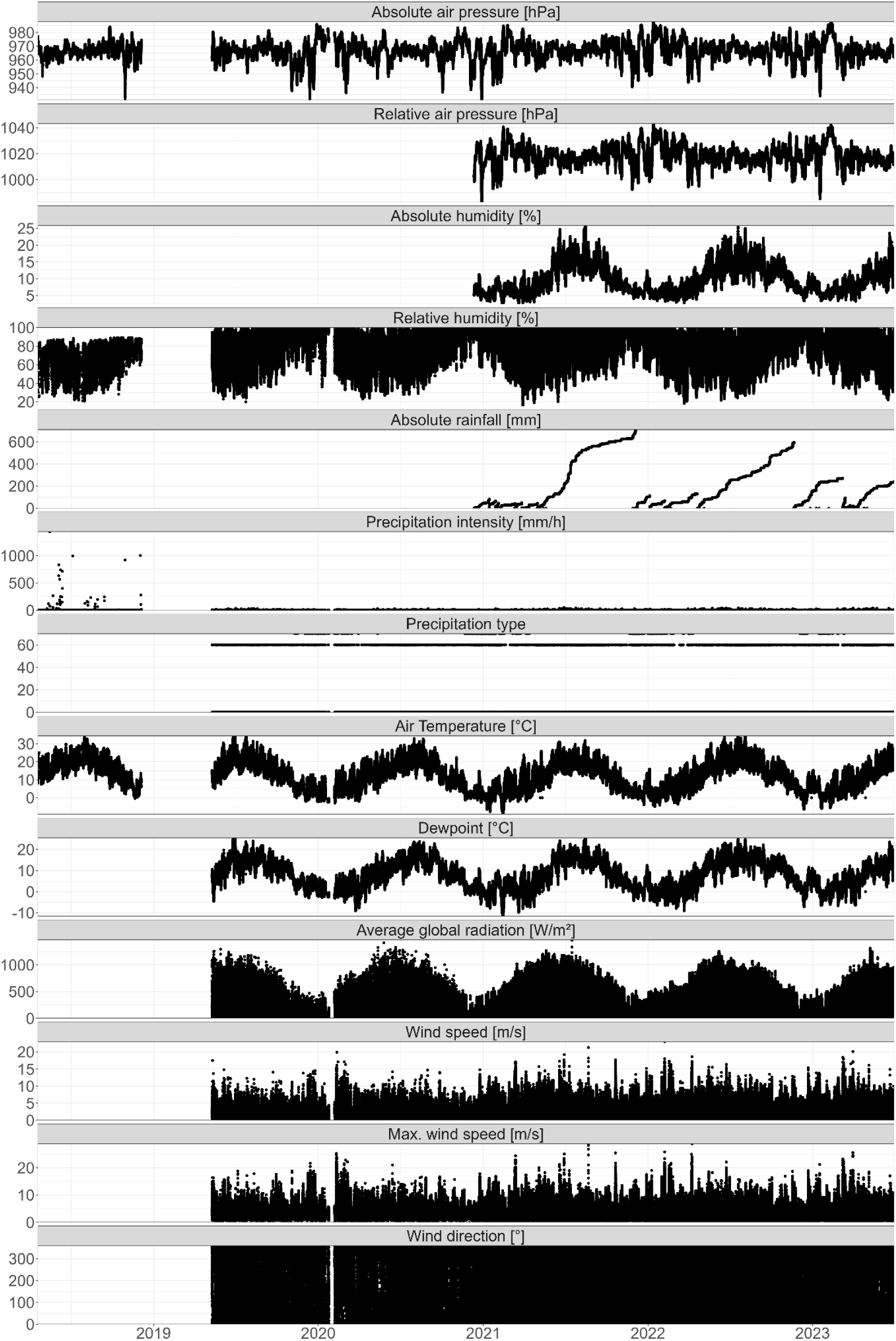
Time series of all meteorological parameters measured from 2018 until June 2023. Each parameter (y-axis) is indicated in the grey box above each panel.

#### Underwater automated dark field imaging microscopy

The instrument with the second highest frequency is the DSPC. Our dual-magnification dark field underwater microscope (DSPC) is based on the Scripps Plankton Camera (SPC) system ^23^. Two magnifications (i.e. two microscope objectives) were designed to cover freshwater plankton imaging in the size range of between ∼ 10 μm - 150 μm for the higher magnification (5.0x objective, with an imaged volume of 0.2 to 10 μL, depending on the depth of field), and between ∼ 100 μm - 1 cm for the lower magnification (0.5x objective, with an imaged volume of 4 to 200 μL). Together, the two magnifications cover the majority of the freshwater plankton food web, offering the largest dynamic size range in the field of underwater plankton imaging ^11^. As an *in-situ* instrument, water flows freely between the two viewports of the camera ^11^. We collected data from 22 May 2018 until 30 June 2023 with a larger gap in the first year from 5 December 2018 to 21 March 2019. With the help of a cronjob (time-based job scheduler software) on the DSPC, the camera system is switched on every hour and images the plankton community for 10 minutes with a frame rate of one frame per second in two magnifications (in 2018 twice a day (starting at 2 am and 2 pm) for 120 minutes). Within each frame, each region of interest (ROI) is detected and isolated from the frame with the use of a Canny edge detector (see Materials and procedures in ^23^ for details of on-board processing) and then saved as a greyscale .tif image (**Fig. 3a**). Each ROI name contains information about the camera and magnification it originated from, as well as the acquisition time in Unix time and further information on the location of the ROI within the original frame (**Fig. 3b**). The raw data is then copied from the internal computer inside the DSPC to the server at Eawag. A Python program is executed hourly on the Eawag server with the help of a scheduled task, which reads in, processes and publishes all newly dated images (see Data Processing). The images are later processed from the Eawag server **(Fig. 3c**, Data Processing Section). Once a week, the camera and its viewports are cleaned.

**Fig. 3:**
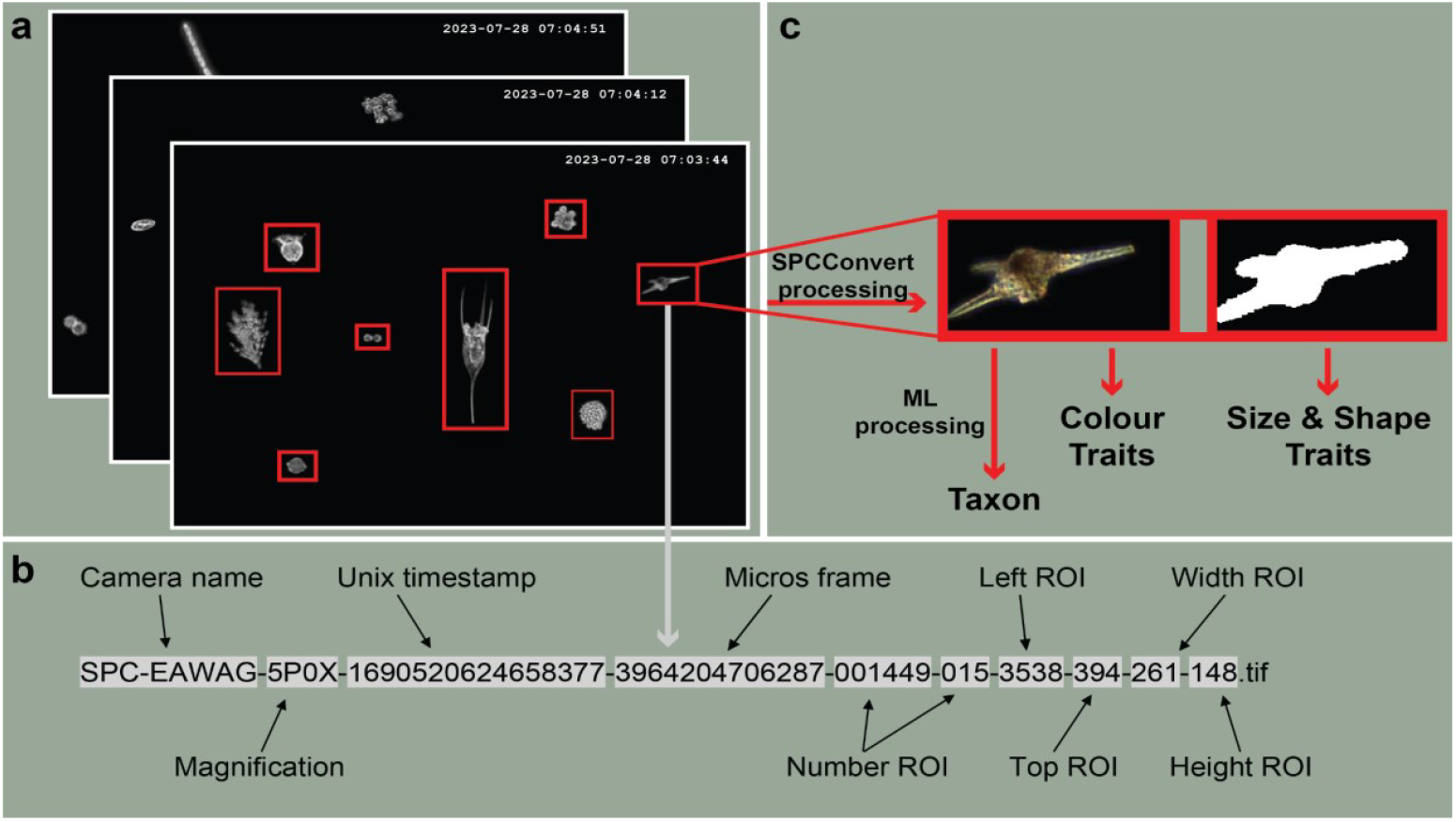
DSPC data processing from whole frames to individual-level trait information and classification. **a)** From each frame, all individual objects are cropped into regions of interest (ROI). **b)** The name of each ROI contains information about the image itself. **c)** Images are processed to extract traits and classification (see Data Processing Section).

#### Automated vertical profiling

The CTD probe used for profiling is an Ocean Seven 316Plus (IDRONAUT S.r.l, Brugherio, Italy) and was changed on 1 July 2020 with the old model being the same as the new one. There is an additional photosynthetically active radiation (PAR) sensor (Li-193 Spherical Underwater Quantum Sensor, LI-COR Environmental, Lincoln, Nebraska) and a fluorimeter (TriLux, Chelsea Technologies Ltd, Surrey, UK). The CTD probe is attached to a winch and installed in the middle below the monitoring platform where it can freely descend and ascend along the water column. The measurement interval is 3 (spring - autumn) - 6 hours (winter). We collected data from 19 April 2018 until 30 June 2023 with a larger gap in the first year from 5 December 2018 to 13 March 2019 where the CTD probe was removed from the monitoring station during the winter months. In January 2022, the PAR and Fluorescence measurements were not taken for four weeks due to the manufacturer servicing of the CTD probe and a temporary replacement with a second machine of the same model that did not fit a PAR sensor and fluorimeter. Gaps in the time series also arise from the removal of measurement errors. In 2020, Phycoerythrin measurements were replaced with Phycocyanin due to the replacement of the TriLux fluorimeter. Before this replacement, the fluorimeter frequently malfunctioned, causing turbidity measurements to fall outside the valid range. Removing these erroneous values created an even larger gap in the turbidity time series.

Most of the year, while physical and biological processes are very dynamic, the measurement frequency was one profile every 3 hours while in winter (usually December - February) when the lake is mixed and power production is low, the profiling frequency was set every 6 hours. The resting position of the CTD probe is at 15 m (in the hypolimnion). For each profile, the winch ascends the probe to 1 m depth, waits 3 minutes to let the water column re-stabilise after the probe recovery, and then lowers it while measuring to a maximum depth of 17 m (2018: part of the year 12 m) with a winch speed of 5 cm/s. During the stratification period, the water column stratification is stable over time until winter mixing, which confirms that the movement of the CTD probe along the water column does not disrupt the stratification of the lake (**Fig. 4**). The parameters are reported every 0.1 m. Further configurations of the CTD probe and profiler can be found in the supplementary information (SI **Tab. S3**). The data is saved in an ASCII format on an Eawag FTP server and automatically converted to online graphics using Python scripts. The graphs are publicly available at https://sensors-eawag.ch/greifensee/main.php. The parameters measured are temperature, conductivity, O_2_ (saturation and concentration), pH, turbidity, PAR, Chlorophyll a, Phycoerythrin and Phycocyanin concentration. In 2020, the measurement of Phycoerythrin was replaced by Phycocyanin (**Fig. 4**). The pH (pH 7 buffer solution) and O_2_ (O_2_ saturation of the atmosphere at 435 masl) sensors are calibrated and the other sensors cleaned once a week (biweekly in winter). In addition to the regular weekly calibrations and cleanings, the CTD probe underwent manufacturer servicing in January 2016. It was replaced with a new but identical probe in July 2020, which subsequently received manufacturer servicing in January 2022.

**Fig. 4:**
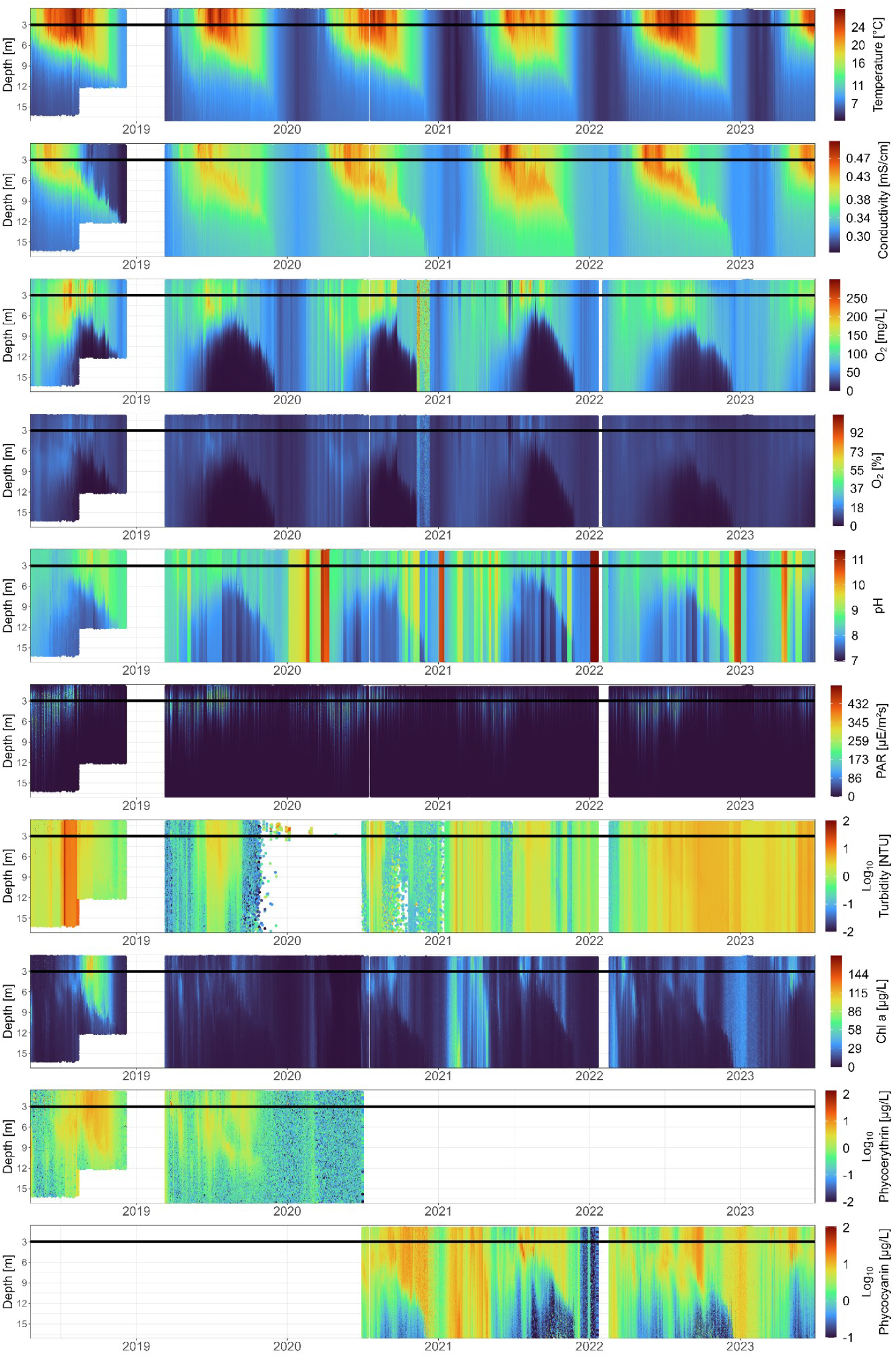
Time series of all CTD parameters measured from 2018 until June 2023. The horizontal line represents the depth at which the DSPC is installed (3 m). For visibility, turbidity, phycoerythrin and phycocyanin are in Log_10_ scale. A non-log figure can be found in SI **Fig. S2**.

#### Nutrient chemistry

We sampled water for nutrient chemistry manually at the monitoring station once per week (or biweekly in winter) from 2 April 2019 to 29 June 2023. In 2018, we did not sample any water for nutrient analysis. We sampled the following nutrients: nitrate (NO_3_ ^-^-N), nitrite (NO_2_ -N), ammonium (NH_4_ ^+^-N), oP (PO_4_ ^3-^-P), TP, TN and TOC. From 19 May 2022 onwards, we also measured silica (H_4_SiO_4_). Nutrients were measured using DIN Standards (German Institute for Standardization, see ^21,24^).

To sample nutrients from discrete depths, we used a 5 L Niskin bottle. Throughout the entire sampling period, we always sampled at 3 m depth (installation depth of the DSPC). We further sampled at 15 m depth (in the hypolimnion) from 27 August 2020 until 29 June 2023. Occasionally, we sampled discrete water samples at depths outside of the regular 3 and 15 m (e.g. at the depth of the Chlorophyll a maximum, **Fig. 5**). For integrated water samples (0 - 18 m), we used a Schröder sampler (2 April 2019 until 12 December 2020). To sample TOC, we used a sediment trap. Sediment trap samples were collected from 25 June 2019 until 2 December 2020 with a sediment trap of 6 cm diameter at 9 m depth (see SI **Fig. S3**).

**Fig. 5:**
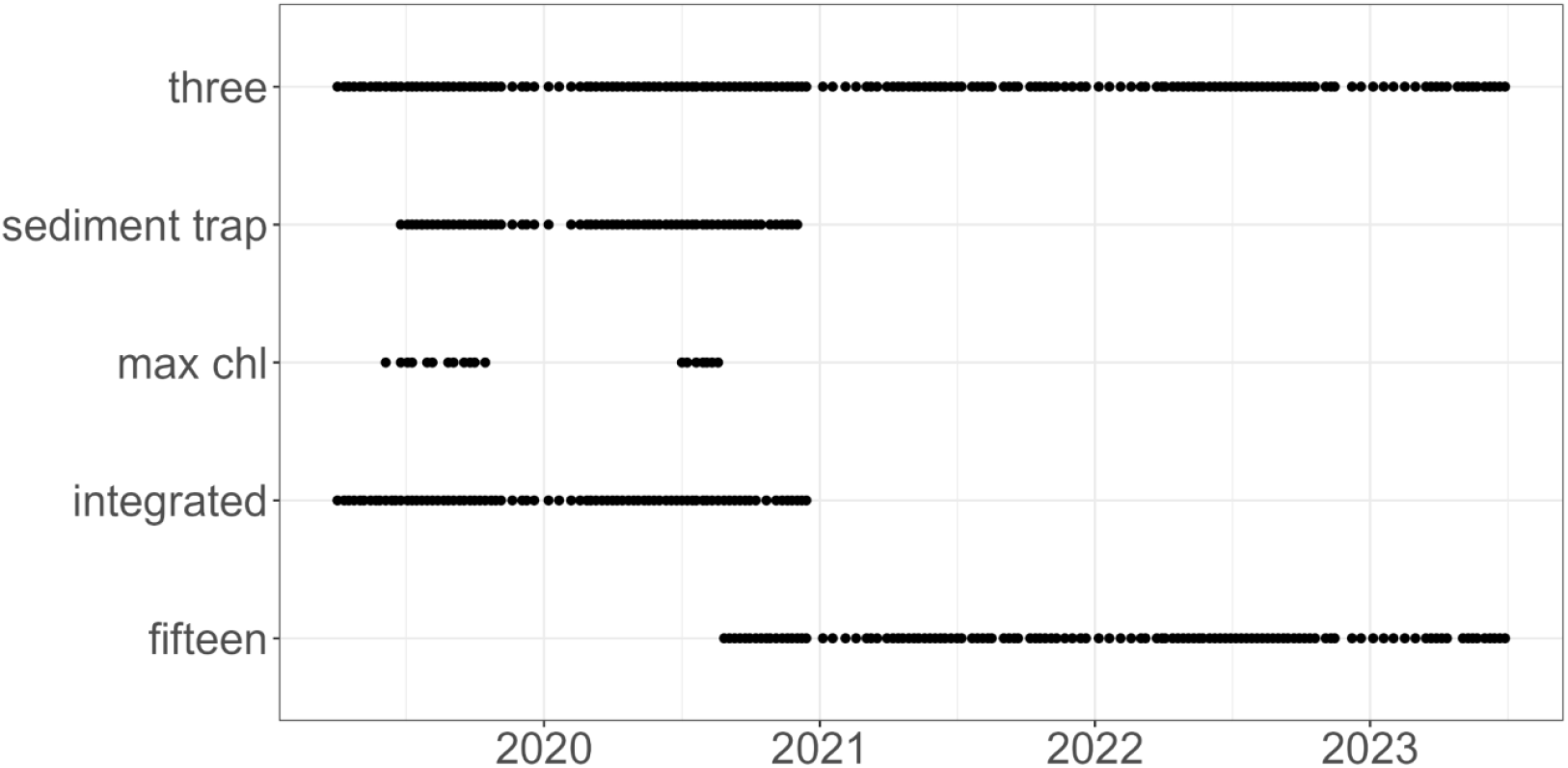
Nutrient chemistry samples taken during the study period 2019 - 2023. Samples were generally taken (bi-)weekly. Here we plot the main samples excluding exceptional samples that do not fall into one of the categories in the plot. Three = 3 m depth, max chl = depth at Chlorophyll a maximum, integrated = 0 - 18 m integrated, fifteen = 15 m depth.

### Data processing

#### Meteorological station

For the meteorological data, we excluded technical artefacts and measurement errors. We removed all time points where every variable recorded a value of 0, as this is a strong indication of malfunctioning of the entire meteo-station. Many variables measured at the meteorological station do not normally include 0 within their expected range, and even for those where 0 is technically possible, such cases are highly unlikely to occur across all variables simultaneously.

Additionally, we set data points where relative humidity (expected range: >0% – 100%) or absolute air pressure (expected range: ∼900 – ∼1050 hPa) equalled 0 to NA, as these values are physically impossible. Since we cannot determine how these errors occurred, we consider all associated measurements at those time points unreliable.

We also set wind measurements (speed and direction) with values of 999.9 to NA, as these are outside the valid ranges (speed: 0 – ∼50 m/s; direction: 0° – 360°) and likely result from technical artefacts, such as electronic noise affecting the sensors.

Finally, we did not correct for sensor drift, as this should be addressed by users in a manner suited to their specific research objectives.

#### Underwater automated dark field imaging microscopy

To process the raw greyscale images (ROI) generated by the DSPC, we use an adapted version of SPCConvert (a collection of Python scripts to convert ROI images into statistics and image mosaics ^25^ developed by Paul L. D. Roberts and Eric Orenstein). We extract colour and binary images from the raw images that are produced by the DSPC (see **Fig. 3c**), as well as individual-level traits ^26^. From the binary images, we extract size and shape traits which are saved in files called features.csv inside the data repository (see the Data Records Section). The traits we extract are the following: major and minor axis length, area, estimated volume, aspect ratio, eccentricity, orientation, and solidity. From the colour images, we extract colour distributions, which information is also found in the files called features.csv. We further feed the colour images to a previously trained machine learning classifier to classify each object/image (ROI) into a taxonomic group ^27,28^. The taxonomic classification for each image is saved in the files called Ensemble_models_Plankiformer_predictions_geo_mean_finetuned.txt inside the data repository (see the Data Records Section). For a list of all taxonomic groups, as well as their taxonomic identity and machine learning accuracy, consult SI **Fig. S4, Fig. S5** and the supplementary table SI_Tab_ML_taxonomy.xlsx. For the lower magnification targeting mesozooplankton and large phytoplankton colonies, we further subsampled the raw image to every 6th second to reduce repeated images of the same object ^11^.

#### Automated vertical profiling

For each profile of the CTD data, we removed depths out of range (0 m - 18 m) and multiple depth measurements of the same depth within a profile (e.g. m 1.1, 1.2, 1.3, 1.3, 1.4, 1.5 […] 15.1, 15.1, 15.1, 15.2 […]) and only kept the first measurement of each depth. These multiple measurements come from the settings of the profiler: we set the CTD probe to acquire sensor readings every 10 cm, but sometimes the probe sends two measures within the same 10 cm range, perhaps due to multiple uncertain readings of the pressure sensor when the profile is disturbed by waves. Further, we removed all measurements from a profile that were taken after reaching that profile’s maximum depth (e.g. m 16.7, 16.8, 16.9, 17, 1.5, 1.6, 1.7, 1.8 […]). This can occur due to a delay in the pressure sensor readings and the controller module stopping the profiling winch. We set measurements that were out of range for a temperate eutrophic lake (i.e. Temperature [°C]: 0 < x < 50; Turbidity [NTU]: 0 < x < 200; Chlorophyll a [μg/L]: 0 < x < 200; Phycocyanin & Phycoerythrin [μg/L]: x > 0, x ≠ 1000; O_2_ [mg/L]: x > 0; O_2_ saturation [%]: x < 300) to NA. We further set O_2_ concentration values where the O_2_ saturation values were below 0 to NA. We did this because the CTD probe internally converts the saturation values into concentrations and we therefore do not trust the concentrations where the saturation was out of range. We further set all negative photosynthetically active radiation (PAR) measurements to 0.

### Data Records

The datasets for abiotic data from the meteorological station, the CTD probe and nutrient measurements are available at the Eawag Research Data Institutional Collection ERIC open (https://doi.org/10.25678/000C8P) ^29^. We provide one data frame (.csv) per instrument and a metadata file (.xlsx) containing variable names and their units.

The biotic datasets from the DSPC are available at the Eawag Research Data Institutional Collection ERIC open (https://doi.org/10.25678/000C2G) ^30^. Each month of our monitoring campaign includes a data package, which consists of one or more compressed files for each magnification and data type: raw data, processed images, and features and classification. The data is organised into folders named with a Unix timestamp for each hour. The raw data includes greyscale .tif images stored in a .tar file, while the processed images are converted to colour .jpeg format after processing with SPCConvert. The features and classification folder contains a features file (features.csv) that details individual size, shape, and colour traits extracted from each ROI through SPCConvert, as well as a classification file (Ensemble_models_Plankiformer_predictions_geo_mean_finetuned.txt) that assigns a taxon to each ROI.

To automate data download from ERIC open, follow the instructions at <u>Automation — ERIC Documentation</u> in the sections “Retrieving information about a package” and “Downloading resources”.

### Technical Validation

#### Meteorological station

We find that data cleaning removed 33.58% of all data points across all variables. The total number of data points left is 16,662,570 across 13 variables and 1,218,393 time points (SI **Fig. S6**).

We compared five years of data that included the change of the meteorological station to a meteorological station by the Federal Office of Meteorology and Climatology MeteoSwiss 7.5 km northwest of the monitoring platform in Fluntern. The MeteoSwiss station is located on the other side of a hill inside the city of Zurich at 556 masl (coordinates: 47° 22’ 40” N, 8° 33’ 56.7” E). We find that while there are local differences between the two meteorological stations, as expected, but the overall patterns between the two stations and locations show the same general patterns (**Fig. 6**).

**Fig. 6:**
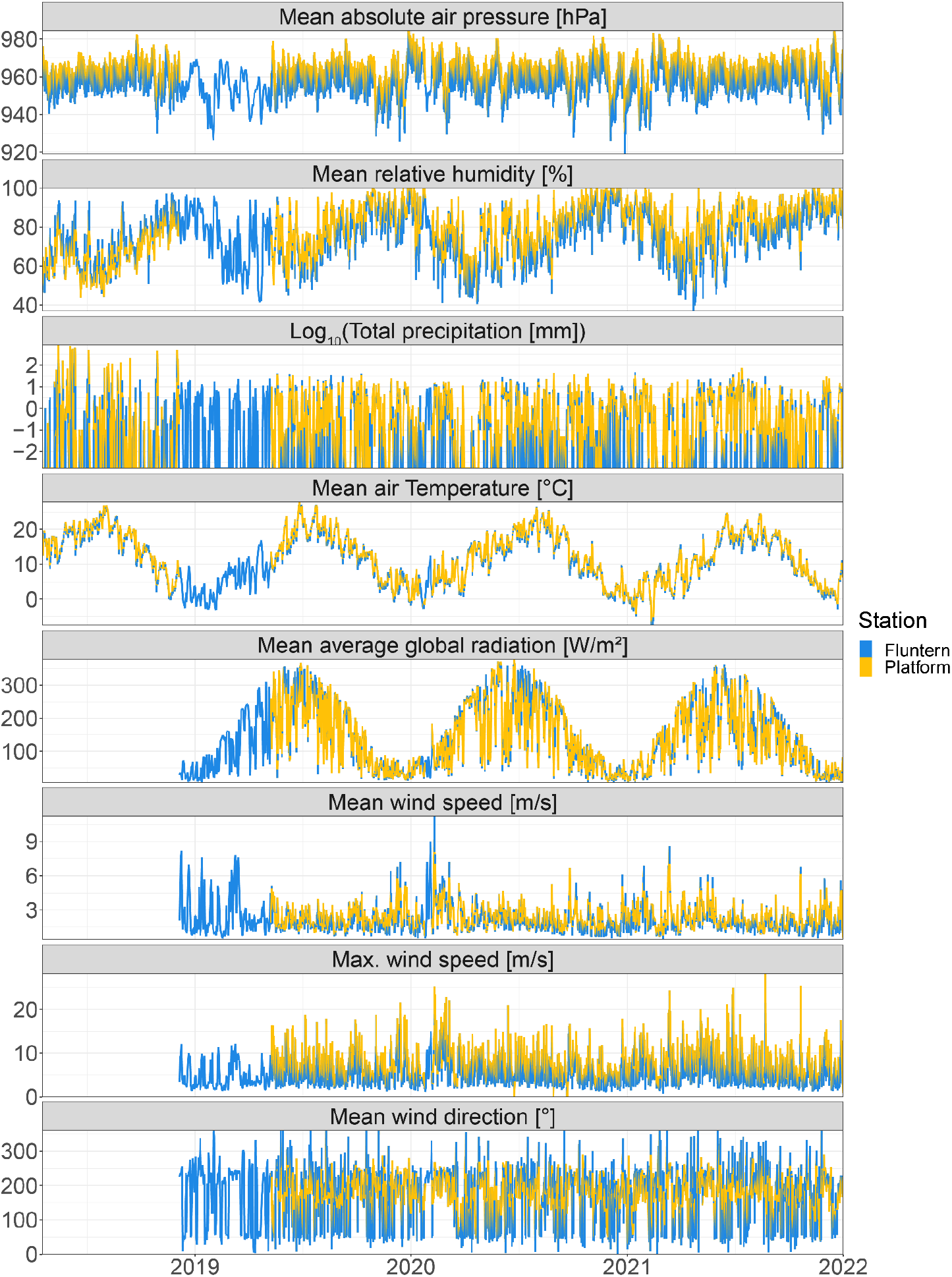
Daily values of a subset of meteorological parameters measured in 2018-2021 at the monitoring platform and the MeteoSwiss station in Fluntern. Each parameter (y-axis) is indicated in the grey box above each panel. Total precipitation at the platform is calculated as the mean precipitation intensity (mm/h) per day multiplied by 24. Note that the total precipitation is in Log_10_ scale.

#### Underwater automated dark field imaging microscopy

In comparison to traditional plankton microscopy data (cells or individuals per litre or m^3^), the data generated by the DSPC comes as ROI per frame or second (in this case the same as we have an imaging frame rate of one frame per second). We have previously shown that ROI/sec is a valid proxy for cell density ^11^. For this, one can divide the total number of ROIs by the number of frames (or seconds) at each measurement time point (folder). Note that for the 0.5x magnification, the number of seconds per hour compared to the 5.0x magnification is reduced to ? as described in the Data Processing Section.

As described in the Methods Section, in 2018, data was acquired twice a day for 120 minutes which resulted in the same amount of minutes as in the rest of the time series where we acquired data 24 times a day for 10 minutes (0.5x magnification with a subset to 1 in 6 seconds). If we aggregate the data to daily observations in ROI/sec, we can see that the time series are comparable across the two acquisition methods in terms of order of magnitude and temporal behaviour **Fig. 7**).

**Fig. 7:**
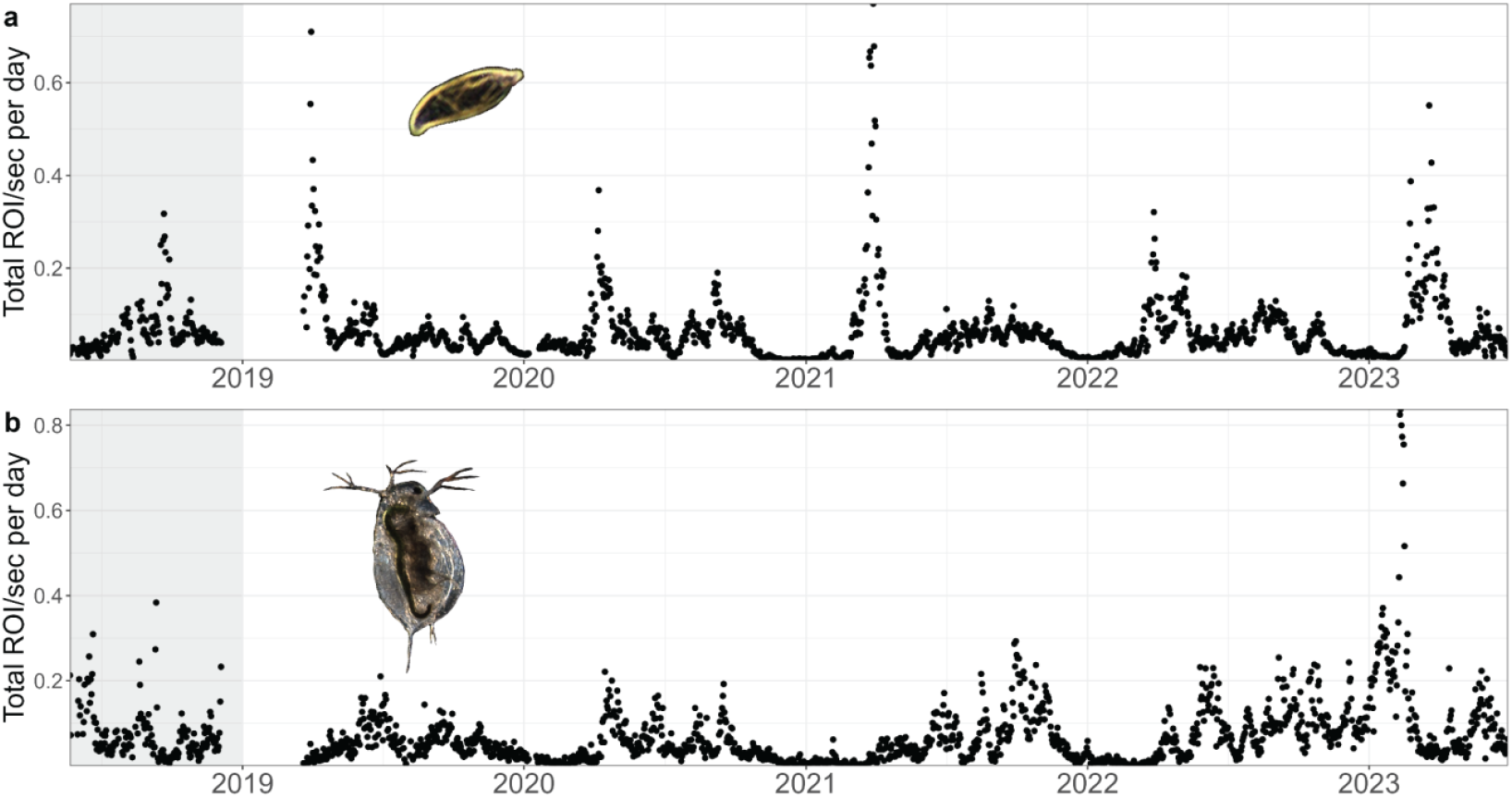
Time series of two common taxa in the **a)** 5.0x magnification (cryptophyceae) and **b)** 0.5x magnification (*Daphnia sp*.). In 2018 (grey area), the distribution of sampling time across the day was different from the rest of the time series.

The relative (and absolute) abundances of taxa are comparable within a magnification. If we want to compare abundances across the magnifications, things are a bit more complex. The theoretical imaging volume of the two magnifications differs by a factor of 20 (5.0x = 0.2 - **10** μL, 0.5x = 4 - **200** μL → **200 μL : 10 μL = factor 20**). When we compare the number of ROI/sec of a taxon that lies within the detection range of both magnifications (*Asterionella sp*.), we can see that the realised difference between the ROI/sec detected in the two magnifications is 22.74 (R^2^ = 0.66, **Fig. 8a**). This number is relatively close to the expected scaling factor of 20.

**Fig. 8:**
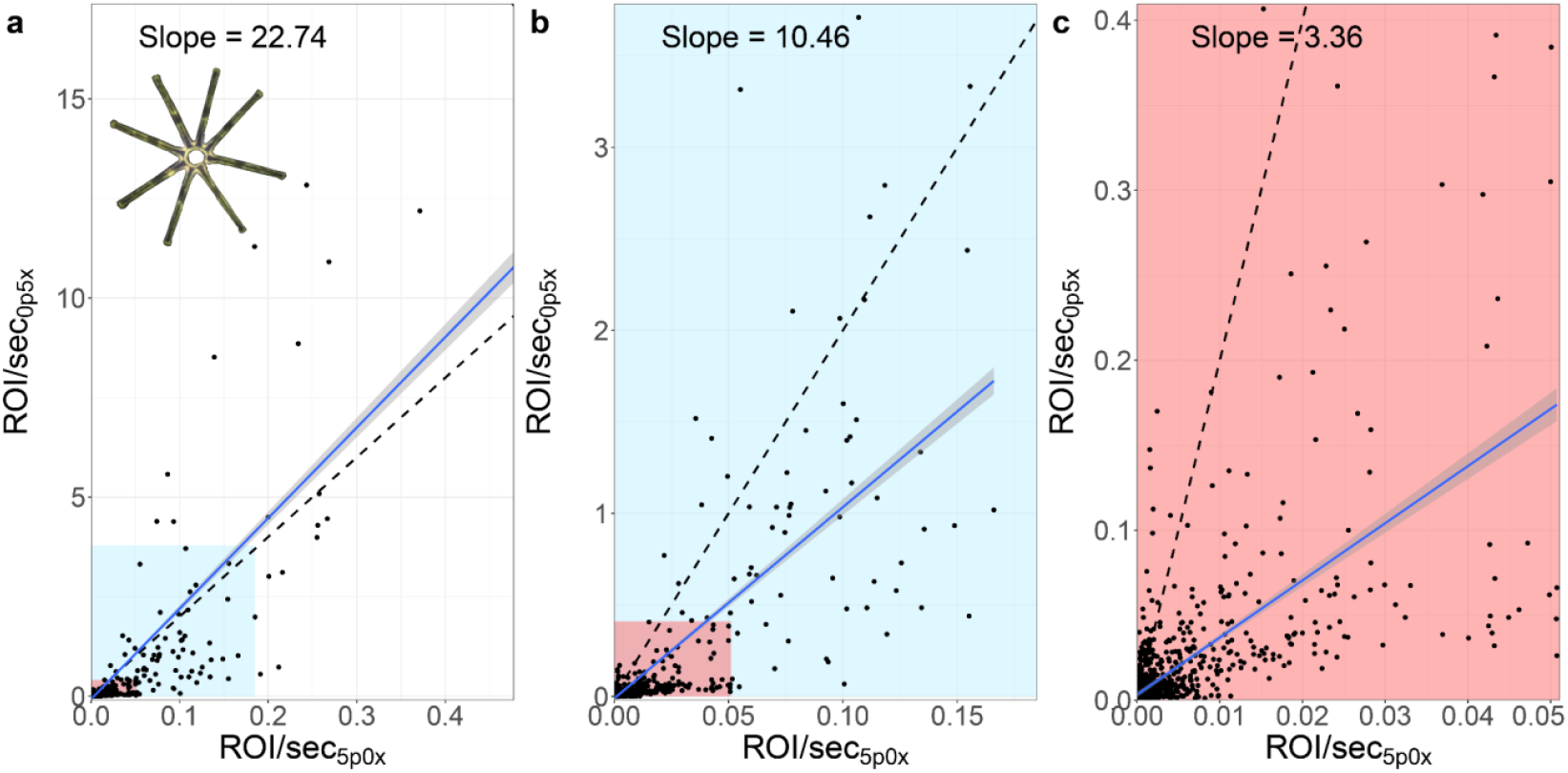
Comparison of ROI/sec from both magnifications on *Asterionella sp*., which has a colony diameter of about 100 μm and lies well within the detection range of both magnifications. The slope of the linear relationship between the two magnifications (daily values 2019 - June 2023) is illustrated as a blue smooth line with the theoretical slope of 20 as a dashed line. **a)** All data points and the 99% and 95% quantiles as coloured blocks, **b)** 99% quantile, **c)** 95% quantile.

Nevertheless, we can see that there are many data points clustered at the lower left corner in **Fig. 8a**. If we exclude the most extreme top 1% (i.e. 99% quantile) of data in both magnifications, the slope decreases to 10.46 (R^2^ = 0.58, **Fig. 8b**) and 3.36 excluding the most extreme top 5% (95% quantile, R^2^ = 0.44, **Fig. 8c**). This discrepancy could arise due to i) turbidity differences affecting the depth of the field of view or ii) the fact that in the 5.0x magnification we capture partial or single colonies in one ROI while in the 0.5x magnification we generally capture single colonies or aggregations of multiple colonies in one ROI.

As an alternative to scaling the two magnifications, one can use the calibration from ROI/sec to individuals/mL from Fig. 3 in ^11^. In theory, after applying the linear scaling factors to each magnification, one can merge the information of the two magnifications. Nevertheless, the calibration was limited to a few species of phytoplankton and *Daphnia* under laboratory conditions ^11^.

We have shown that as of now, the data generated by a DSPC in the laboratory does not fully match traditional microscopy data after using the calibration from ^11^. We argue that this might be a limitation due to the limited species pool in the calibration excluding large colonies and motile species ^31^. We therefore recommend using that approach with caution and keeping its limitations in mind.

Further limitations of the data generated by the DSPC presented here include i) a limitation to the detection of translucent objects (very dense colonies (e.g. *Woronichinia sp*.) or inorganic particles (e.g. microplastics) are not detected with the DSPC) and ii) only objects with a size above ∼10 μm, iii) no detection of surface phytoplankton blooms due to its position at 3 m depth or iv) the underestimation of species richness/diversity as illustrated in Fig. 5 in ^11^.

#### Automated vertical profiling

We find that data cleaning removed 14.12% of all data points across all variables. The total data points left is 14,068,236 across 10 variables and 10,517 time points (SI **Fig. S7**).

We compared three years of data to monthly measurements taken by the Zurich Cantonal authorities 2 km south of our monitoring station, at the deepest point of the lake (coordinates: 47° 20′ 59.54″ N 8° 40′ 41.13″ E, SI **Fig. S1**). While there are local differences between the two measurements, the overall patterns between the two stations and locations correlate (**Fig. 9**). We cannot benchmark all parameters measured by the CTD probe against traditional monitoring, but this comparison, as well as the wide use of CTD probes across the globe and its sensors (https://gleon.org/), confirms that automated CTD profiling is a robust approach to water physical and chemical parameter monitoring.

**Fig. 9:**
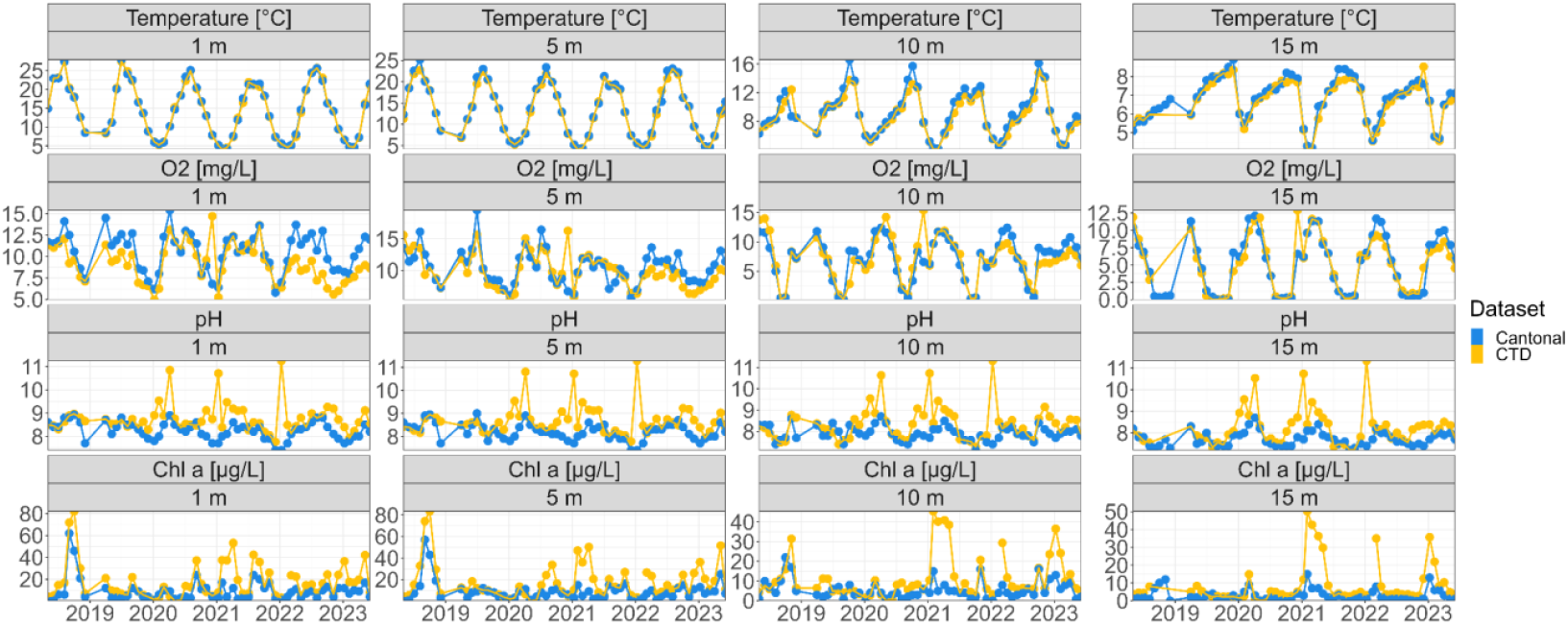
Daily values of a subset of physicochemical parameters measured in 2018-2023 at the monitoring platform by the CTD probe and monthly values measured by cantonal authorities. Each parameter (y-axis) at different depths is indicated in the grey box above each panel.

### Usage Notes

#### General considerations

The dataset presented here provides minimally processed environmental and plankton monitoring data to maximise its utility across diverse research applications. We removed only clear technical artefacts and instrument-related measurement errors, preserving the underlying variability in the raw signals. This curation approach maintains data integrity while avoiding preprocessing steps (such as drift correction, denoising, or outlier removal) that could bias specific analytical applications. The preservation of raw data characteristics is particularly valuable for:

- Ecological researchers investigating plankton abundance patterns or morphological trait dynamics, which may require different temporal or spatial aggregation approaches
- Signal processing specialists developing new methods for environmental sensor data analysis
- Computer vision and machine learning researchers working with plankton imagery
- Aquatic ecosystem modellers and limnologists requiring high-resolution data for time series analysis and model calibration

By providing data in this minimally processed form, researchers can implement processing workflows optimised for their specific research questions, whether focusing on data science, high-frequency ecological dynamics, or modelling developments.

#### Underwater automated dark field imaging microscopy

As described in the Methods Section, the measurement frequency and interval differed in 2018 compared to the rest of the time series. Theoretically, each measurement time point (folder) should consist of 120 minutes of data acquisition. This is not always the case. We therefore recommend incorporating the difference between the acquisition time of the first and the last ROI of each folder to get a better estimate of the total number of seconds (frames) to calculate the ROI/sec. We further recommend making sure that there is only one folder for each hour (in UTC) for the time series 2019 - 2023 as multiple folders suggest a technical error.

As (zoo-)plankton is known to exhibit diel vertical migration ^32^ we recommend aggregating data to daily observations. While we have already shown that zooplankton in Lake Greifen shows a pattern of diel vertical migration ^11^, some phytoplankton have been shown to exhibit similar patterns (^33^ and SI **Fig. S8)** which supports this recommendation. To aggregate, we further recommend setting a threshold of a minimum number of hours of data acquisition to have unbiased and comparable results across the time series.

#### Automated vertical profiling

First, we want to address the issue of photosynthetically active radiation (PAR) measurements under the monitoring station. As the CTD profiler ascends and descends from the centre of the monitoring platform, the first part of the profile is shaded. This results in an increase in measured PAR values with depth to a maximum of PAR. After reaching the maximum PAR, intensities decrease with depth as is generally expected (**Fig. 10**). The amount of shading depends on multiple factors and is not constant (**Fig. 10b**). The depth and intensity of shading most likely depends, among other things, on the incoming global radiation (affected by cloud cover), the water’s turbidity and the time of day and year.

**Fig. 10:**
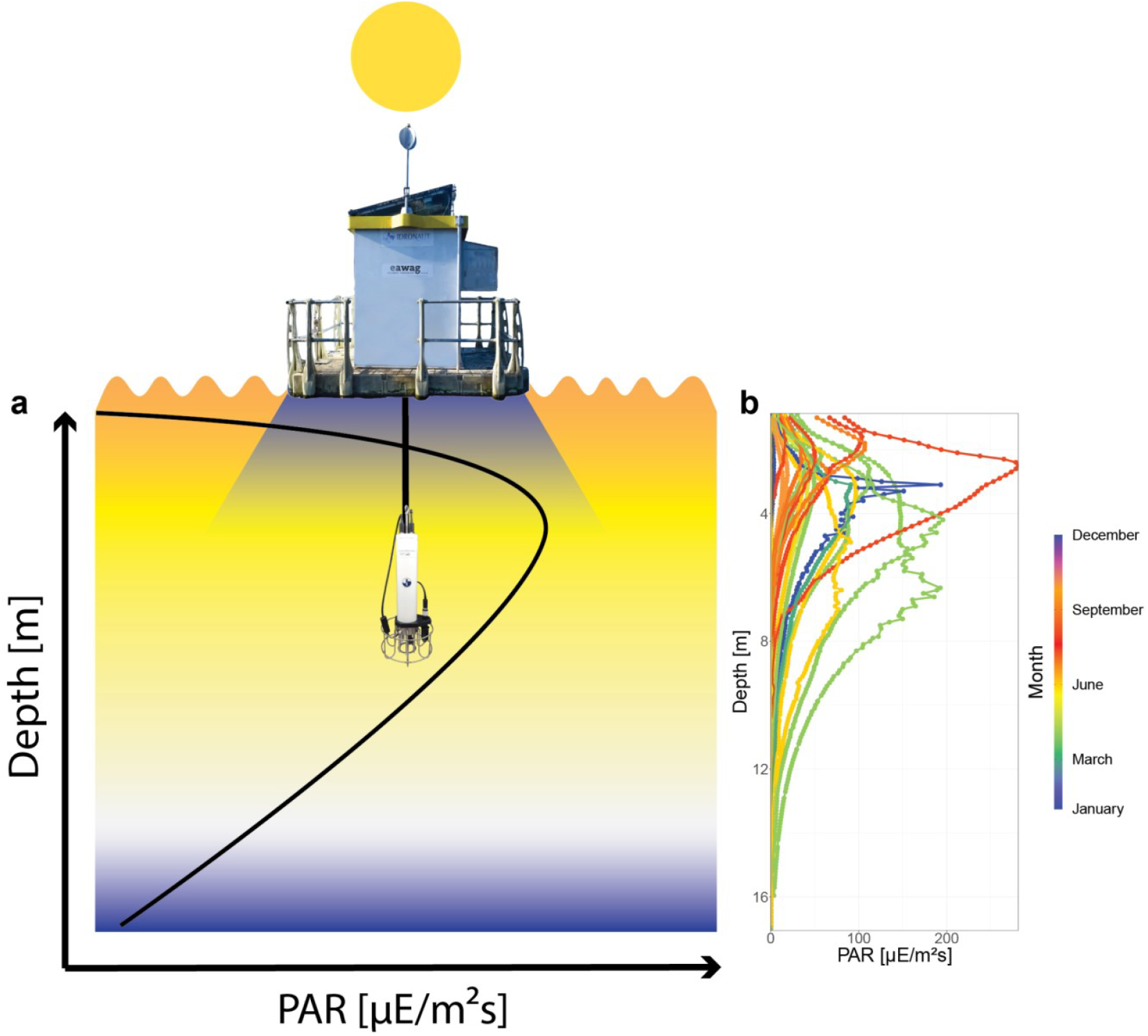
PAR profiles by the CTD probe at the monitoring platform. **a)** Schematic illustration of the CTD setup below the platform and its influence on PAR measurements. The CTD probe descends below the platform and is therefore shaded in the first few metres. **b)** 75 profiles across the 5 years and the influence of the month on the shape of the depth-PAR curve.

As shown in the pH depth profiles (**Fig. 4, Fig. 9**), high pH values are observed at the monitoring platform during winter. This pattern is consistent across multiple years, indicating a recurring physical or chemical process rather than a sensor malfunction. In winter, pH remains relatively uniform throughout the water column, which is expected when the lake fully mixes during turnover (see other parameters measured by the CTD probe, **Fig. 4**). Note that pH is only calibrated biweekly during winter (Dec-Feb). In addition, the pH readings might be influenced by conductivity and temperature. The lake has a reduced buffering capacity at lower conductivity, making pH more variable. At low temperatures, pH sensors take longer to stabilise, which may affect reliability ^34^. In Lake Greifen, winter temperatures drop to 3.5°C, and conductivity can be as low as 0.3 mS/cm. While these factors may contribute to the observed pH values, they do not necessarily indicate faulty data across the time series. We encourage readers to consider these aspects in their analysis, as accounting for potential sensor drift or environmental influences may refine the interpretation of pH trends.

Finally, even after cleaning data from doubled measurements at the same depth and removing measurements after the profile is finished, you will still find profiles of different lengths (depth ranges). There are several profiles where the maximum depth (∼17 m) is not reached. This should be taken into account when aggregating data from different depths and time points.

#### Nutrient chemistry

Note that when measurements are below the detection limit (specified by the method and analysis device - SI **Tab. S4**), this is indicated by a less-than sign (<) and the detection limit for a given specific nutrient. We generally recommend replacing that number with a small number close to or clearly under the detection limit.

Further, when using the TOC values from the sediment trap, it should also be noted that although most sampling dates are one week apart, the exact number of days can vary. Occasionally the sampling dates were shifted by a few days or the samples were taken biweekly. When comparing the TOC values over the time series, the interval between the sampling dates must be taken into account.

## Supporting information

SI_Tab_ML_taxonomy

## Code Availability

The script to convert raw images into colour images and individual-level traits can be found at ^26^. For the classification of images, the code can be found at ^27^. The scripts that were used to further analyse data from our monitoring data can be found in past ^21,35,36^ and in future publications from Eawag. Raw data processing scripts for the meteorological station and the CTD probe can be found in the supplementary information (SI **Tab. S5, Tab. S6**).

## Acknowledgement

This research was funded by the Swiss National Science Foundation (projects 182124 Aquascope and 202290 Cyanobloom) and the German Research Foundation (project 412375259 Aquascope), with contribution for infrastructure by the Swiss Federal Office for the Environment (contract Nr Q392-1149). F.P. and M.B.-J. acknowledge the Eawag DF project Big-Data Workflow (#5221.00492.999.01).

## Author Contributions

Conceptualization of the monitoring program and manuscript was done by FP, MR, M B-J, CE and SD. Funding acquisition was done by FP and M B-J. SE and SD were responsible for data curation. SE, EM and PN contributed to the formal analysis. Methodology was done by SE, MR, FP and M B-J. SE, MR, FP and M B-J were responsible for project administration. The software was the responsibility of CE, SD and M B-J. Validation was done by EM, SE, M B-J, MR and PN. Visualisation by SE and EM. The original draft was written by SE and FP. All authors contributed to the review and editing of the manuscript.

## References

1. Williamson, C. E., Saros, J. E. & Schindler, D. W. Climate change. Sentinels of change. Science 323, 887–888 (2009). doi:10.1126/science.1169443.

2. Carrer, M., Dibona, R., Prendin, A. L. & Brunetti, M. Recent waning snowpack in the Alps is unprecedented in the last six centuries. Nat. Clim. Chang. (2023) doi:10.1038/s41558-022-01575-3.

3. Jasechko, S. et al. Rapid groundwater decline and some cases of recovery in aquifers globally. Nature 625, 715–721 (2024). doi:10.1038/s41586-023-06879-8.

4. Xu, F.-L., Tao, S., Dawson, R. W., Li, P.-G. & Cao, J. Lake ecosystem health assessment: Indicators and methods. Water Res. 35, 3157–3167 (2001). doi:10.1016/s0043-1354(01)00040-9.

5. Kallis, G. The EU water framework directive: measures and implications. Water Policy 3, 125–142 (2001). doi:10.1016/s1366-7017(01)00007-1.

6. Kane, D. D., Gordon, S. I., Munawar, M., Charlton, M. N. & Culver, D. A. The Planktonic Index of Biotic Integrity (P-IBI): An approach for assessing lake ecosystem health. Ecol. Indic. 9, 1234–1247 (2009). doi:10.1016/j.ecolind.2009.03.014.

7. Lyche Solheim, A. et al. Ecological threshold responses in European lakes and their applicability for the Water Framework Directive (WFD) implementation: synthesis of lakes results from the REBECCA project. Aquat. Ecol. 42, 317–334 (2008). doi:10.1007/s10452-008-9188-5.

8. Stockwell, J. D. et al. Storm impacts on phytoplankton community dynamics in lakes. Glob. Chang. Biol. 26, 2756–2784 (2020). doi:10.1111/gcb.15033.

9. Anneville, O. et al. The paradox of re-oligotrophication: the role of bottom–up versus top–down controls on the phytoplankton community. Oikos 128, 1666–1677 (2019). doi:10.1111/oik.06399.

10. Rogers, T. L. et al. Trophic control changes with season and nutrient loading in lakes. Ecol. Lett. 23, 1287–1297 (2020). doi:10.1111/ele.13532.

11. Merz, E. et al. Underwater dual-magnification imaging for automated lake plankton monitoring. Water Res. 203, 117524 (2021). doi:10.1016/j.watres.2021.117524.

12. Álvarez, E., Moyano, M., López-Urrutia, Á., Nogueira, E. & Scharek, R. Routine determination of plankton community composition and size structure: a comparison between FlowCAM and light microscopy. J. Plankton Res. 36, 170–184 (2013). doi:10.1093/plankt/fbt069.

13. Rivas-Villar, D., Rouco, J., Penedo, M. G., Carballeira, R. & Novo, J. Automatic Detection of Freshwater Phytoplankton Specimens in Conventional Microscopy Images. Sensors 20, (2020). doi: 10.3390/s20226704.

14. Rivas-Villar, D., Rouco, J., Carballeira, R., Penedo, M. G. & Novo, J. Fully automatic detection and classification of phytoplankton specimens in digital microscopy images. Comput. Methods Programs Biomed. 200, 105923 (2021). doi:10.1016/j.cmpb.2020.105923.

15. Pomati, F., Jokela, J., Simona, M., Veronesi, M. & Ibelings, B. W. An automated platform for phytoplankton ecology and aquatic ecosystem monitoring. Environ. Sci. Technol. 45, 9658–9665 (2011). doi:10.1021/es201934n.

16. Merz, E. et al. Disruption of ecological networks in lakes by climate change and nutrient fluctuations. Nat. Clim. Chang. 13, 389–396 (2023). doi:10.1038/s41558-023-01615-6.

17. Behrenfeld, M. J. & Boss, E. S. Student’s tutorial on bloom hypotheses in the context of phytoplankton annual cycles. Glob. Chang. Biol. 24, 55–77 (2018). doi:10.1111/gcb.13858.

18. Arteaga, L. A., Boss, E., Behrenfeld, M. J., Westberry, T. K. & Sarmiento, J. L. Seasonal modulation of phytoplankton biomass in the Southern Ocean. Nat. Commun. 11, 5364 (2020). doi: 10.1038/s41467-020-19157-2.

19. Isles, P. D. F. & Pomati, F. An operational framework for defining and forecasting phytoplankton blooms. Front. Ecol. Environ. 19, 443–450 (2021). doi:10.1002/fee.2376.

20. Bürgi, H. R., Bührer, H. & Keller, B. Long-term changes in functional properties and biodiversity of plankton in Lake Greifensee (Switzerland) in response to phosphorus reduction. Aquat. Ecosyst. Health Manag. 6, 147–158 (2003). doi:10.1080/14634980301471.

21. Thomas, M. K., Fontana, S., Reyes, M., Kehoe, M. & Pomati, F. The predictability of a lake phytoplankton community, over time-scales of hours to years. Ecol. Lett. 21, 619–628 (2018). doi:10.1111/ele.12927.

22. Istvánovics, V. Eutrophication of lakes and reservoirs. Lake Ecosystem Ecology; Elsevier: San Diego, CA, USA 47–55 (2010).

23. Orenstein, E. C. et al. The Scripps Plankton Camera system: A framework and platform for in situ microscopy. Limnol. Oceanogr. Methods 18, 681–695 (2020). doi:10.1002/lom3.10394.

24. Public Health Association, A. Standard methods for the examination of water and wastewater. 6, (1926).

25. SPCConvert. (Github). https://github.com/tooploox/SPCConvert

26. SPCConvert: All Versions of SPCConvert, That We Use for Processing of Acquired Images from the Underwater Microscope. (Github). https://github.com/phytoecoeawag/SPCConvert.

27. Kyathanahally, S. P. Plankiformer. (Github). https://github.com/kspruthviraj/Plankiformer.

28. Chen, C. et al. Producing plankton classifiers that are robust to dataset shift. Limnol. Oceanogr. Methods 23, 39–66 (2025). doi:10.1002/lom3.10659.

29. Merkli, S., Reyes, M., Ebi, C., Dennis, S. R. & Pomati, F. Greifensee meteo-CTD-chemistry data 2018 - 2023. Preprint at 10.25678/000C8P (2024).

30. Dennis, S. et al. Aquascope: Raw data, images and classifications - May 2018 to June 2023. Preprint at 10.25678/000C2G (2023).

31. Merkli, S., Reyes, M. & Pomati, F. Laboratory application of the Aquascope approach of automated imaging and classification for long-term plankton monitoring. bioRxiv 2024.02.23.581739 (2024) doi:10.1101/2024.02.23.581739.

32. Bandara, K., Varpe, Ø., Wijewardene, L., Tverberg, V. & Eiane, K. Two hundred years of zooplankton vertical migration research. Biol. Rev. Camb. Philos. Soc. 96, 1547–1589 (2021). doi:10.1111/brv.12715.

33. Olli, K. Diel vertical migration of phytoplankton and heterotrophic flagellates in the Gulf of Riga. J. Mar. Syst. 23, 145–163 (1999). doi:10.1016/S0924-7963(99)00055-X.

34. Plummer, N. & Peck, D. L. pH measurements of low-conductivity waters. Water-Resources Investigations Report (1987) doi:10.3133/WRI874060.

35. Merkli, S. et al. Automated plankton monitoring suggests a key role of microzooplankton and temperature for predicting dynamics of phytoplankton size classes. (2024) doi:10.1101/2024.02.23.581723.

36. Daugaard, U., Merkli, S., Merz, E., Pomati, F. & Petchey, O. L. The dependence of forecasts on sampling frequency as a guide to optimizing monitoring in community ecology. Ecosphere 15, (2024). doi:10.1002/ecs2.4786.

37. Latz, M. A. C. et al. A comprehensive dataset on spatiotemporal variation of microbial plankton communities in the Baltic Sea. Sci Data 11, 18 (2024). doi:10.1038/s41597-023-02825-5.

