## Supplementary_information for "Five years of high-frequency data of phytoplankton zooplankton and limnology from a temperate eutrophic lake"

### Table of content

**Supplementary Figures**

**Fig. S1:** Topography map of Lake Greifen. 2

**Fig. S2:** Non-log time series of all CTD parameters 3

**Fig. S3:** Sediment trap used for TOC measurements. 4

**Fig. S4:** All taxonomic categories of the 0.5x magnification of the DSPC. 5

**Fig. S5:** All taxonomic categories of the 5.0x magnification of the DSPC. 6

**Fig. S6:** Histograms of every parameter in meteorological data. 7

**Fig. S7:** Histograms of every parameter in CTD data. 8

**Fig. S8:** Abundance of the cyanobacteria *Aphanizomenon sp.* across the day. 9

**Supplementary Tables**

**Tab. S1**: Topography table of Lake Greifen. 10

**Tab S2**: Measurement interval and time range of all instruments installed at the monitoring platform in Greifensee. 11

**Tab. S3:** CTD and profiler configurations 12

**Tab. S4:** Detection limits of nutrients according to the analysis device. 16

**Tab. S5:** R-Script used to treat the raw data from the meteorological stations. 17

**Tab. S6:** R-Script used to treat the raw data from the CTD probe. 19

**References** 21

**Supplementary Figures**

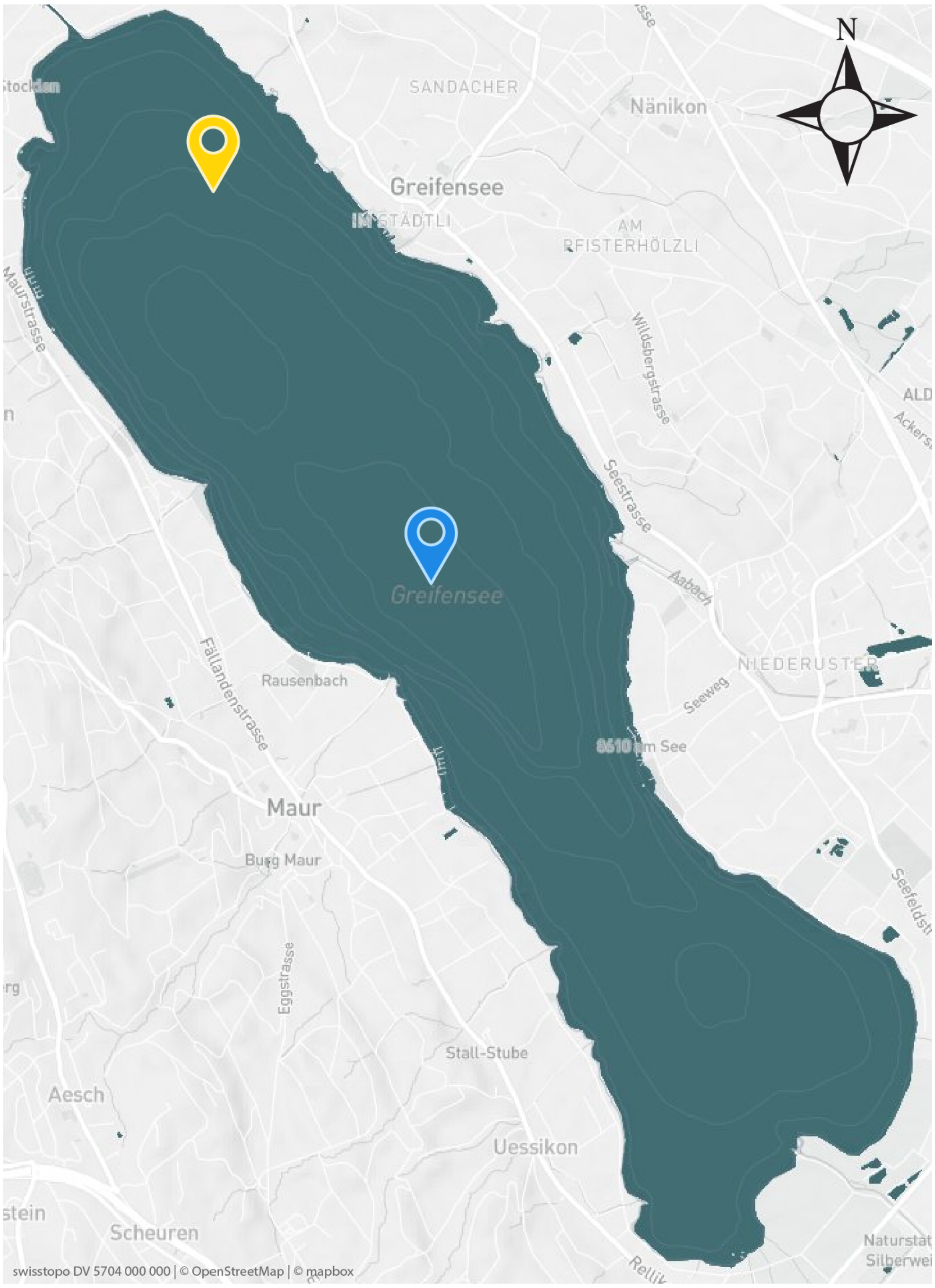


**Fig. S1:** Topography map of Lake Greifen. The map was retrieved from https://www.datalakes-eawag.ch/data. The yellow (northern) pin indicates the location of the monitoring station and the blue (southern) pin indicates the location of the lake's deepest point at 32 m depth.

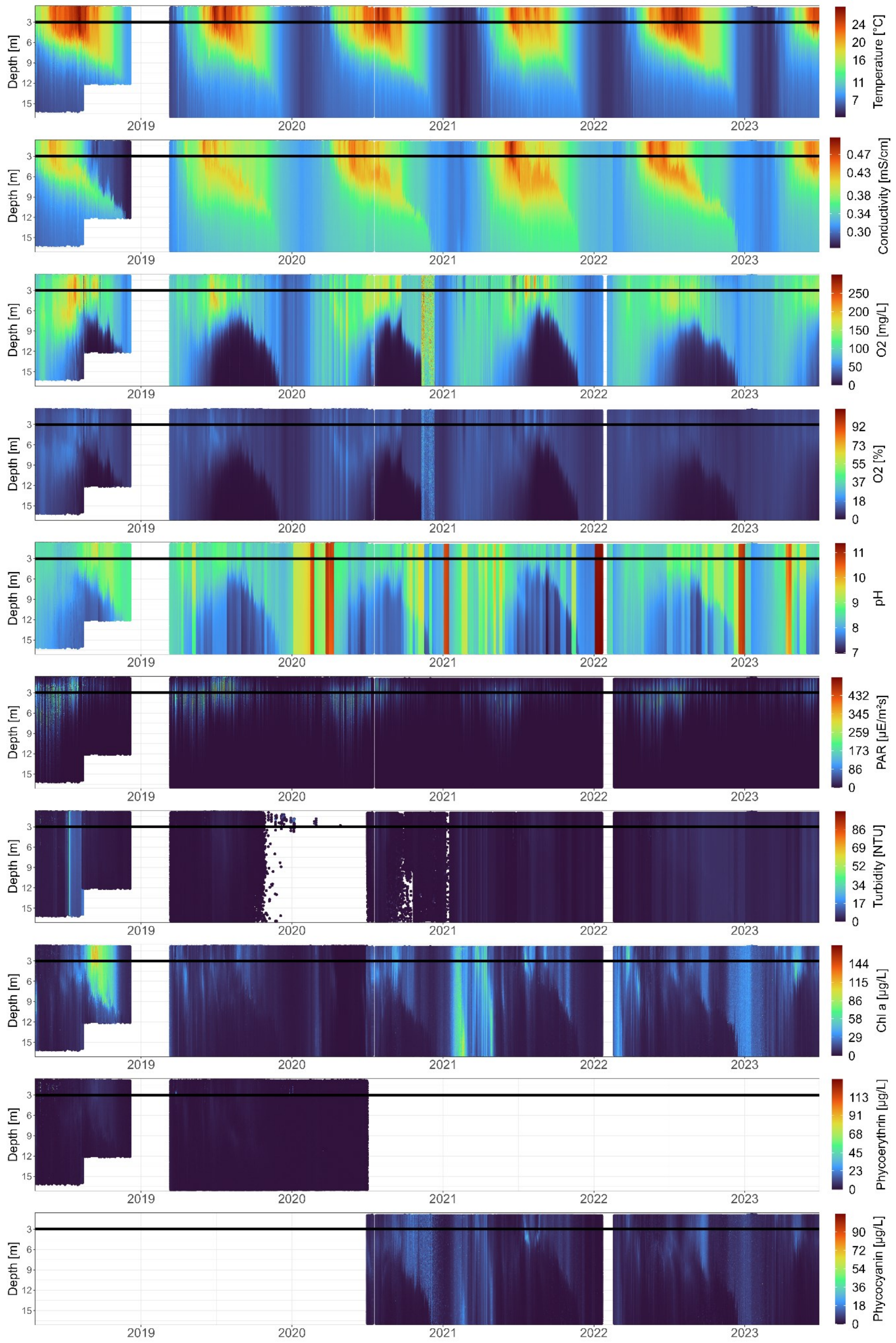


**Fig. S2:** Non-log time series of all CTD parameters measured from 2018 until June 2023. The horizontal line represents the depth at which the DSPC is installed (3 m).

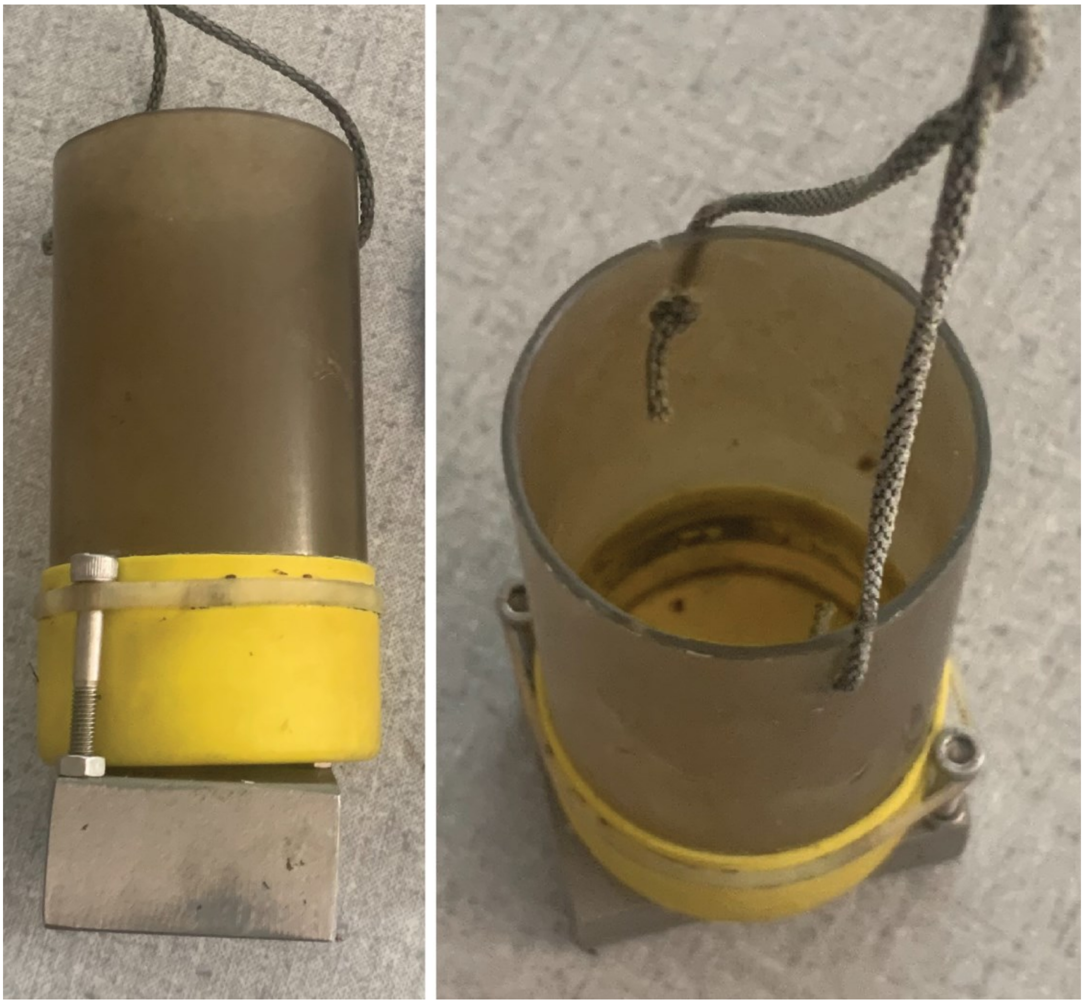

**Fig. S3**: Sediment trap used for TOC measurements from 25 June 2019 until 2 December 2020. 6 cm diameter. Traps accumulated sedimenting particles between sampling dates and were retrieved for nutrient analysis at the sampling date.

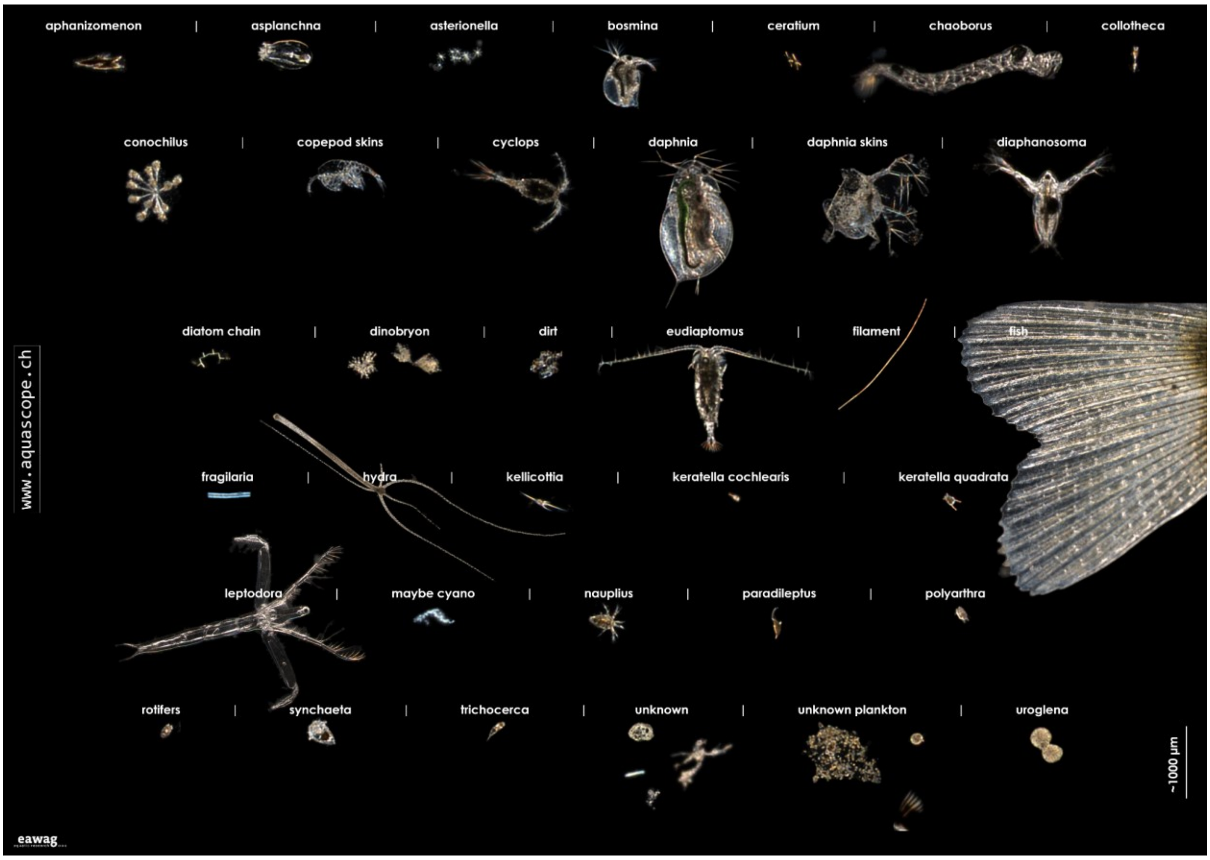


**Fig. S4:** All taxonomic categories generated by the machine learning classifier for the 0.5x magnification of the DSPC.

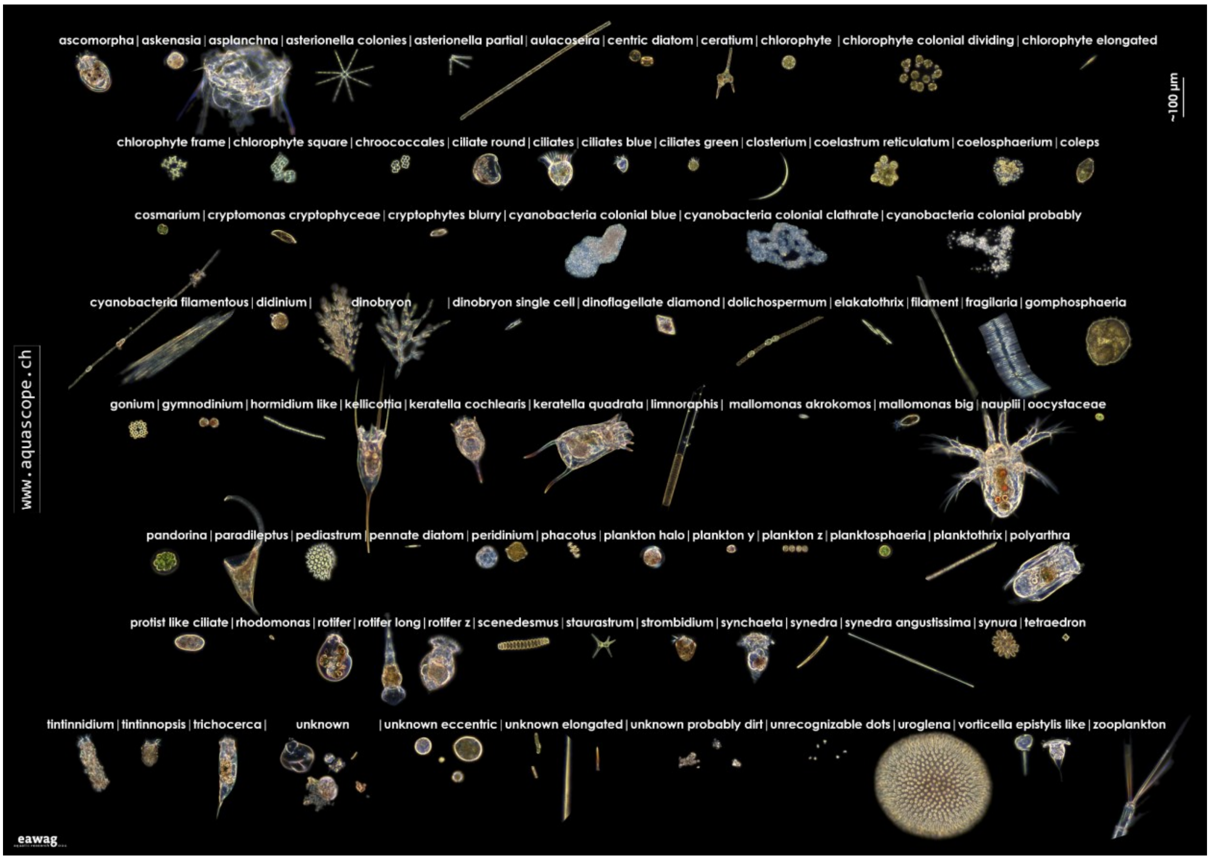


**Fig. S5:** All taxonomic categories generated by the machine learning classifier for the 5.0x magnification of the DSPC.

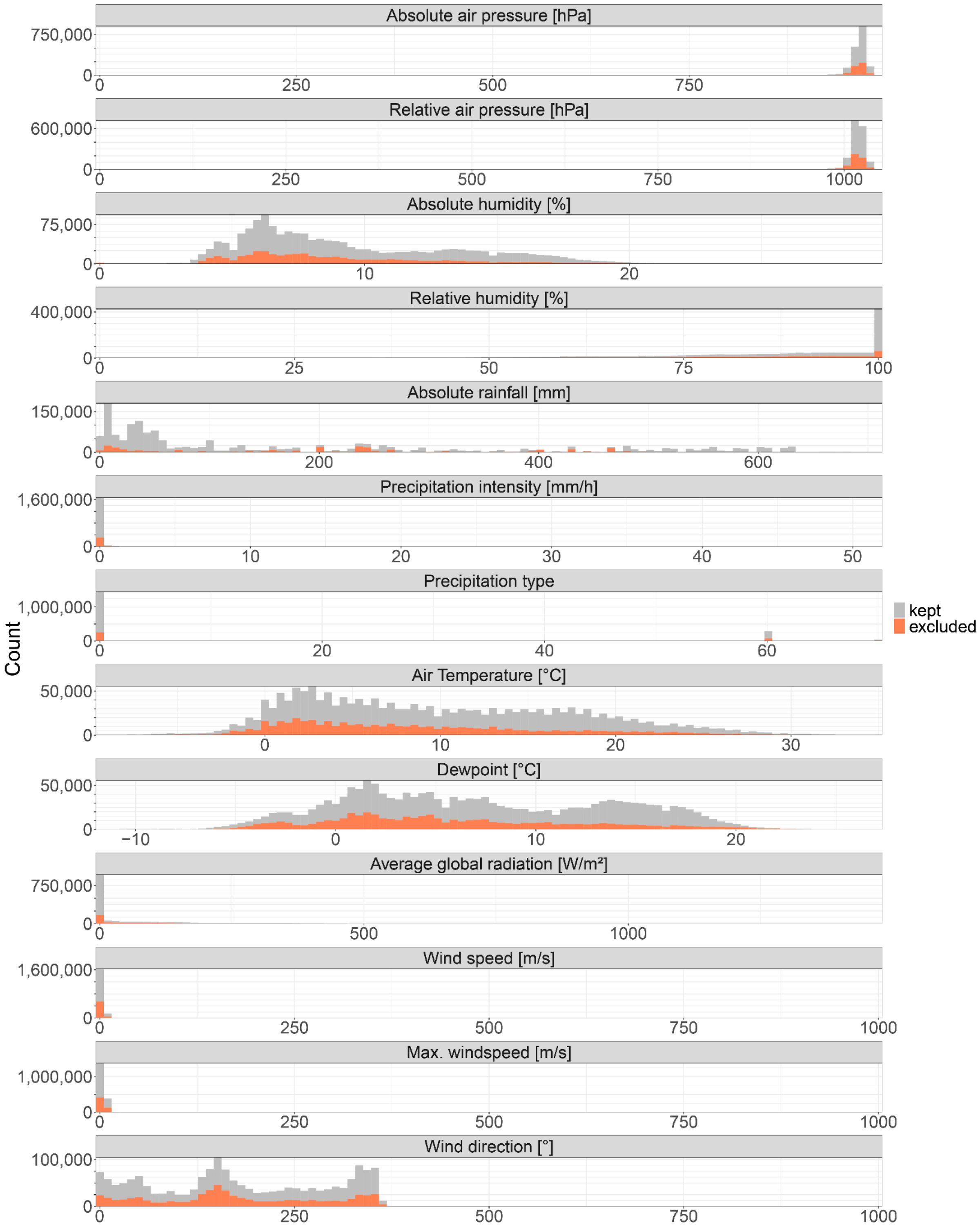


**Fig. S6**: Histograms of every parameter in meteorological data. Counts are displayed on the y-axis and the values of each variable are on the x-axis. In colour, we display the data points excluded in the time series.

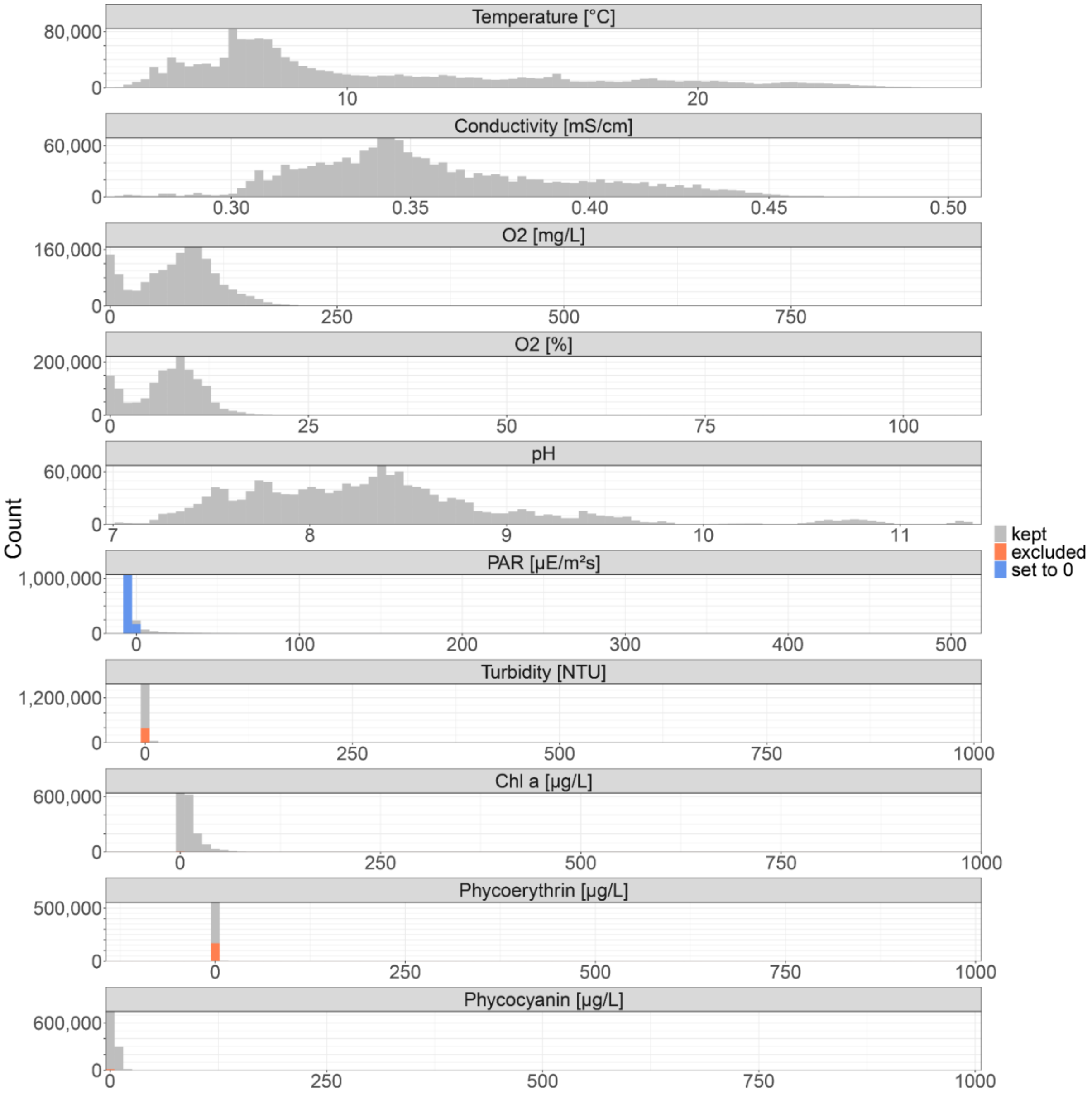


**Fig. S7**: Histograms of every parameter in CTD data. Counts are displayed on the y-axis and the values of each variable on the x-axis. In colour, we display the data points excluded in the time series or set to zero (PAR measurements only).

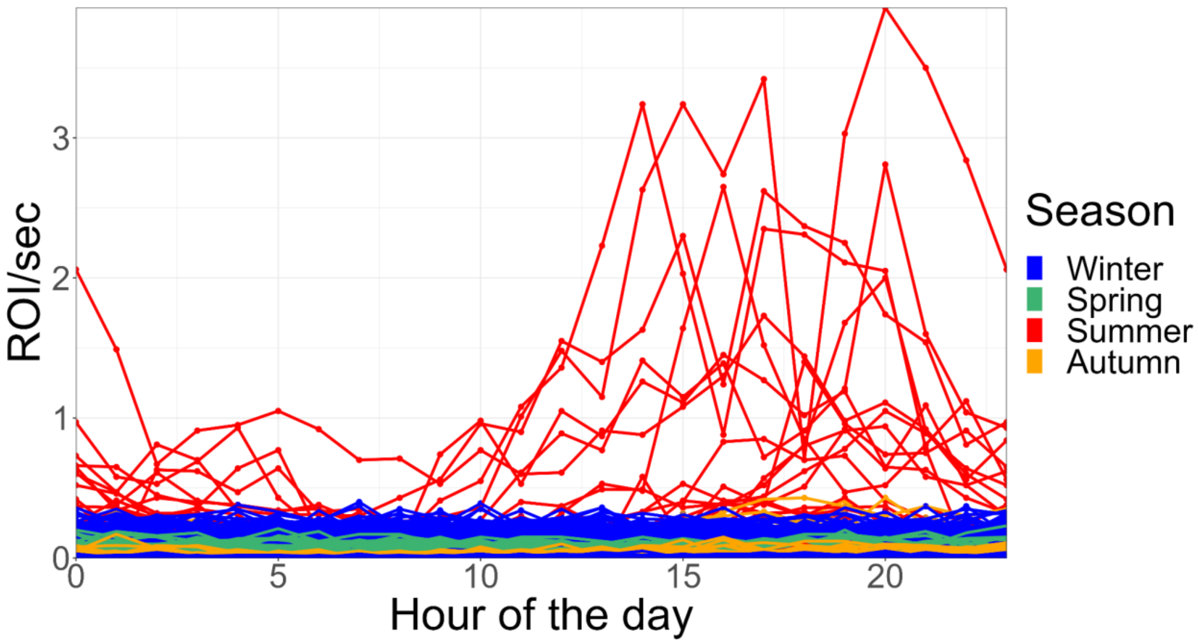


**Fig S8**: Abundance of the cyanobacteria *Aphanizomenon sp.* captured with the 0.5x magnification from 2019 to 2023. Each ROI represents a colony and not a single cell. Each line connects all hours of one day. The colours represent the season (Winter = Dec - Feb, Spring = Mar - Mai, Summer = Jun - Aug, Autum = Sept - Nov). In summer when we find the highest abundance of *Aphanizomenon sp.*, we see a peak of abundance in the late afternoon with lower abundances at night. This suggests a vertical migration of this cyanobacteria species that is known to be able to regulate its buoyancy and migrate vertically across the water column [1].

### Supplementary tables

**Tab. S1**: Topography table of Lake Greifen. For the calculation of e.g. mixed layer depth from temperature profiles, one needs the following information. Here we report the area for each depth from the surface to the maximum depth of 32 m.

| Depth [m] | Area [$km^2$] |
|---|---|
| 0 | 8.447586 |
| 0.3 | 8.403289 |
| 1 | 8.300381 |
| 2.5 | 8.082 |
| 5 | 7.747 |
| 7.5 | 7.177 |
| 10 | 6.467 |
| 12.5 | 5.564 |
| 15 | 4.62 |
| 17.5 | 3.925 |
| 20 | 3.458 |
| 22.5 | 2.999 |
| 25 | 2.468 |
| 27.5 | 1.918 |
| 30 | 0.92 |
| 31 | 0.298646 |
| 32 | 0.017904 |
| 32.3 | 0.0001 |

**Tab S2**: Measurement interval and time range of all instruments installed at the monitoring platform in Greifensee. The end date of the datasets of all instruments is 30.06.2023.  Dates in dd.mm.yyyy format. Note that the measurement intervals noted are the theoretical regular intervals. There might be deviations to those intervals.

| Instrument | Instrument name | Time zone of timestamp | Start date | Data gap winter 2018/2019 | Measurement interval |
|---|---|---|---|---|---|
| Meteorological station | Vaisala Oyj WXT520 | UTC | 18.04.2018 | 05.12.2018 - 10.05.2019 | 30 min |
| | LUFFT WS700-UMB + Meteobridge PRO data logger | UTC | 10.05.2019 | - | 10 min |
| | LUFFT WS700-UMB + netDL 1000 data logger | UTC | 11.12.2020 | - | 1 min |
| DSPC | DSPC | Unix timestamp (UTC) | 22.05.2018 | 05.12.2018 - 21.03.2019 | 1 h |
| CTD | Idronaut Ocean Seven 316Plus | CE(S)T | 19.04.2018 | 05.12.2018 - 13.03.2019 | 3 - 6 h |
| Nutrients | - | CE(S)T | 02.04.2019 | - | (Bi-) weekly |

**Tab. S3**: CTD and profiler configurations used at the monitoring platform. The time between acquisition changes from 3 h to 6 h in winter. The rest of the configurations stay the same all year.

```
Login:..OCEAN SEVEN Probe communication, flush them
StandBy
$$
Present time Sun Nov 26 21:17:40.0 2023
Next timeout Sun Nov 26 21:19:40.0 2023
##

Login:..Verbose mode access to the CMD ShellPassWord <<**********
USER ACCESS
BCUy2k ID:[09.01.00.00]{USR}(1.8_00-01/2020) Sun Nov 26 21:18:49.8 2023
OPERATOR MODE - <HELP>Show the commands list
BCUy2k-Cmd>stop
BCUy2k ID:[09.01.00.00]{USR}(1.8_00-01/2020) Sun Nov 26 21:18:51.4 2023
STOP the automatic monitoring cycle

Do you want to stop the monitoring cycle ?:Yes
Systems status[Done]
STD Memory[Cast000%-Data001%]
EXT Memory[Cast025%-Data011%]
Configuration updating...Done
BCUy2k-Cmd>auxc
BCUy2k ID:[09.01.00.00]{USR}(1.8_00-01/2020) Sun Nov 26 21:19:11.1 2023
Auxiliary systems configuration
AUX SYS:(0)Quit,(1)GSM,(2)METEO,(3)GPS/GALVAPOT,(4)OCEAN SEVEN,(5)ACOUSTIC
METER,(6)Analog
AUX SYS:(7)U-SOUND-CYCLEPO4,(8)RDI-ADCP,(9)Cytosense-VIP-MPCP
Aux. Sys to configure [0..9]:0 enter new value< 4
Do you want to UNINSTALL the OCEAN SEVEN Auxiliary system ?:No
BCUy2k ID:[09.01.00.00]{USR}(1.8_00-01/2020) Sun Nov 26 21:19:37.5 2023
OCEAN SEVEN profiler setup

Probe Code :3
Measuring unit OCEAN SEVEN
Probe:<0>None,<1>OS303,<2>OS304,<3>OS316/320,<4>OS305

Probe type:3
Probe WarmUp       [s]:30
Watch for the data [s]:1
Min. operating current [mA]:1
Max. operating current [mA]:350
Profiler data acquisition mode:<0>Profile,<1>Stationary,<2>Continuous profile

Selection:0
Time between acquisitions or profiles starting from 5-minute
Present timeout: 03:00:00 enter time [hh:mm:ss]  < 06:00:00
Winch speed                  [cm/s]:5
Maximum profiling depth       [dbar]:17
Profiling depth step          [dbar]:0.1
Starting depth offset         [dbar]:1
Sensor package stand-By depth [dbar]:15
Microprofile starting depth   [dbar]:0
```

```
First measure (surface) equilibrium timeout [s]:180
Waiting timeout on acquisition point [s]:0
Depth to judge the validity of the UpperMost probe position[dbar]:1
Type <any key>To continue
--- Configure the parameter to acquire ---
Please, declare the parameters in the same order used by the acquisition system
to transmitt them
IndexID.CodeNameDigitsPrecision
    1000   Press0701
    2001    Temp0803
    3089    Cond0801
    4003  Cond200801
    5106  O2%opt0802
    6107   O2ppm0801
    7007      pH0803
    8025     PAR0701
    9215     CHL0701
   10216    TURB0701
   11213  Phycya0701

CMD:(I)nitialize,(A)dd,(D)elete,(M)odify,(Q)uit
--- Configure the parameter to acquire ---
Please, declare the parameters in the same order used by the acquisition system
to transmitt them
IndexID.CodeNameDigitsPrecision
    1000   Press0701
    2001    Temp0803
    3089    Cond0801
    4003  Cond200801
    5106  O2%opt0802
    6107   O2ppm0801
    7007      pH0803
    8025     PAR0701
    9215     CHL0701
   10216    TURB0701
   11213  Phycya0701

CMD:(I)nitialize,(A)dd,(D)elete,(M)odify,(Q)uit
Number of bytes used per acquisition 48
Type <any key>To continue
BCUy2k ID:[09.01.00.00]{USR}(1.8_00-01/2020) Sun Nov 26 21:23:05.8 2023
Auxiliary systems configuration
AUX SYS:(0)Quit,(1)GSM,(2)METEO,(3)GPS/GALVAPOT,(4)OCEAN SEVEN,(5)ACOUSTIC
METER,(6)Analog
AUX SYS:(7)U-SOUND-CYCLEPO4,(8)RDI-ADCP,(9)Cytosense-VIP-MPCP
Aux. Sys to configure [0..9]:0
Configuration updating...Done
BCUy2k-Cmd>0
StandBy
$$
Present time Sun Nov 26 21:23:40.0 2023
Next timeout Sun Nov 26 21:25:40.0 2023
##
```

```
Login:..Verbose mode access to the CMD Shell
BCUy2k ID:[09.01.00.00]{USR}(1.8_00-01/2020) Sun Nov 26 21:23:43.0 2023
OPERATOR MODE - <HELP>Show the commands list
BCUy2k-Cmd>
BCUy2k-Cmd>strt
BCUy2k ID:[09.01.00.00]{USR}(1.8_00-01/2020) Sun Nov 26 21:24:02.0 2023
Start an automatic monitoring cycle

Warning the monitoring starting time must be greater than the present time

Monitoring starting time
Current value: 21:24:02 enter time [hh:mm:ss]  < 00:00:00
Warning the monitoring starting time must be greater than the present time

Monitoring starting time
Current value: 21:24:08
BCUy2k-Cmd>240   star
 !!Error in command code
BCUy2k-Cmd>t strt
BCUy2k ID:[09.01.00.00]{USR}(1.8_00-01/2020) Sun Nov 26 21:24:55.5 2023
Start an automatic monitoring cycle

Warning the monitoring starting time must be greater than the present time

Monitoring starting time
Current value: 21:24:55 enter time [hh:mm:ss]  < 23:59:00
Do you want to clear the RunTime status informations?:Yes
BCU y2k automatic monitoring switch-OFF in progress..
Systems status[Done]
STD Memory[Cast000%-Data001%]
EXT Memory[Cast025%-Data011%]
OS3xx   - Next Acq. Sun Nov 26 23:59:00.1 2023

STD Memory[Cast000%-Data001%]
EXT Memory[Cast025%-Data011%]
Configuration updating...Done
Present time Sun Nov 26 21:25:11.3 2023
Next timeout Sun Nov 26 23:59:00.3 2023

Wait at least 10 seconds before executing a new Start-Upx€x€€xx€x€xxxøxø€xxxøxxx€xøø

BCUy2k-Id:[09.01.00.00](1.8_00-01/2020) -Sun Nov 26 21:25:33.0 2023
HW Setup...done
Memory...[Cnf.oK][Fw.oK][Rt.oK][Data.oK][Ext.00495 MByte Verify.oK]
STD Memory[Cast000%-Data001%]
EXT Memory[Cast025%-Data011%]

WarmUp......
Analogue...Battery(12.124 V) Solar Panel Not installed
ComPorts...GSM.OS
RunTime...GSM.OS3xx.ANA..RTC PowerOn, Start a Measurement Cycle
BCUy2k ID:[09.01.00.00]{USR}(1.8_00-01/2020) Sun Nov 26 21:25:40.6 2023
Automatic monitoring startup
```

Systems status[Done]
STD Memory[Cast000%-Data001%]
EXT Memory[Cast025%-Data011%]UMTS COMMUNICATIONS

UMTS Comms-WarmUp-

56

**Tab. S4:** Detection limits of nutrients according to the analysis device.

| Parameter | Detection limit | Analysis device |
|---|---|---|
| Nitrate | 0.1 | Metrohm 930 Compact IC Flex |
| Nitrite | 1 | Spektrophotometer Agilent Cary 60 |
| Ammonium | 5 | |
| oP | 1 | |
| TP | 3 | |
| TN | 0.5 | Shimadzu TOC-L CSH |
| TOC | 0.5 | |
| Silica | 0.5 | Skalar San++ Autoanalyzer |

**Tab. S5**: R-Script used to treat the raw data from the meteorological stations. Cleaning steps for both meteorological stations in BLACK. Cleaning steps for Vaisala Oyj WXT520 in **BLUE**. Cleaning for LUFFT WS700-UMB in **GREEN**.

```
# import metadata of manually merged variable names

metadata_meteostations <- read_excel("~/3 Data Management Projects/10 the data maker aka
chocolate factory/METEO/Meteostation_Greifensee_metadata_vaisala_OTTStations.xlsx",
                                   sheet = "meteostations_variablelist")
metadata_meteostations <- dplyr::select(metadata_meteostations, -unit)

#  Vaisala Oyj WXT520

metadata_meteostations_old <- dplyr::filter(metadata_meteostations, !is.na(vaisala_old_variable))

meteodata_vaisala <- read_delim("MeteoGreifensee.csv",
                     delim = ";", escape_double = FALSE, trim_ws = TRUE)
# remove timestamp column (first of the two date variables):
# seems to be a unlogic one as it has the same value for multiple entries (2019-05-15 11:41:00) 649
times all other only once.
# DateTime has one value for each observation only
meteodata_vaisala <- dplyr::select(meteodata_vaisala, -c(ID, timestamp)) # also get rid of ID as we
don't need that
meteodata_vaisala$DateTime <- as.POSIXct(as.character(meteodata_vaisala$DateTime), format =
"%d.%m.%Y %H:%M")

# rename the variables according to the metadata
for(i in 1:ncol(meteodata_vaisala)){
  if(colnames(meteodata_vaisala)[i] %in% metadata_meteostations_old$vaisala_old_variable){
    colnames(meteodata_vaisala)[i] <-
metadata_meteostations_old$final_variable_name[metadata_meteostations_old$vaisala_old_variab
le == colnames(meteodata_vaisala)[i]]
  }
}

# use only the variables that are in the metadata
meteodata_vaisala <- meteodata_vaisala[,c(metadata_meteostations_old$final_variable_name)]


# LUFFT WS700-UMB

# load data
meteodata <- read_delim("meteo_greifensee.csv",
                     delim = ";", escape_double = FALSE, trim_ws = TRUE)

# delete variables we don't need + make date a posixct
meteodata_lufft  <- dplyr::select(meteodata_lufft, -c(id, stationId, name,  batteryVoltage,
gsmSignal))
meteodata_lufft$timestamp <-  as.POSIXct(as.numeric(meteodata_lufft$TST), origin="1970-01-
01") # as.POSIXct(as.character(meteodata_lufft$timestamp), format = "%Y-%m-%d
%H:%M:%OS")
meteodata_lufft <- dplyr::select(meteodata_lufft, -TST)

# change the column name according to the metadata
```

```
# check if column names are with 0 in front
for(i in 2:ncol(meteodata_lufft)){
  if(substr(colnames(meteodata_lufft)[i],1,1) == 0){
    colnames(meteodata_lufft)[i] <- substr(colnames(meteodata_lufft)[i], 2,
nchar(colnames(meteodata_lufft)[i]))
  } else {
    print(colnames(meteodata_lufft)[i])
  }
}
# rename
metadata_meteostations_new_name <- dplyr::filter(metadata_meteostations,
!is.na(OTTStation_new_id))
for(i in 1:ncol(meteodata_lufft)){
  if(colnames(meteodata_lufft)[i] %in% metadata_meteostations_new_name$OTTStation_new_id){
    colnames(meteodata_lufft)[i] <-
metadata_meteostations_new_name$final_variable_name[metadata_meteostations_new_name$OT
TStation_new_id == colnames(meteodata_lufft)[i]]
  }
}

# use only the variables that are in the metadata
meteodata_lufft <- meteodata_lufft[,c(metadata_meteostations_new_name$final_variable_name)]

# cleaning data across all data

meteo_final <- bind_rows(meteodata_vaisala, meteodata_lufft)

# there are some observations that have 0s everywhere -> kill those observations
meteo_final_cleaned <- meteo_final[!rowSums(meteo_final[, 2:9])==0,]

# also likely erroneous data: windspeed = 999.9 -> kill observations by replacing them with NA
(only wind varibales are affected)
meteo_final_cleaned$windspeed[meteo_final_cleaned$windspeed == 999.9] <- NA
meteo_final_cleaned$winddirection[meteo_final_cleaned$winddirection == 999.9] <- NA
meteo_final_cleaned$windspeed_max[meteo_final_cleaned$windspeed_max == 999.9] <- NA

# zeros in humidity and pressure make are impossible values -> set all those values to NA
meteo_final_cleaned$rownr <- seq(1:nrow(meteo_final_cleaned))
meteo_final_cleaned_zeros <- dplyr::filter(meteo_final_cleaned, rel_humidity == 0 |
abs_airpresssure == 0)
# set all values of that measurement to NA
meteo_final_cleaned_zeros[,2:15] <- NA
# re-join and sort again
meteo_final_cleaned_new <- bind_rows(meteo_final_cleaned_nozeros,
meteo_final_cleaned_zeros)
meteo_final_cleaned_new <- meteo_final_cleaned_new[ order(meteo_final_cleaned_new$rownr ,
decreasing = F ),]
meteo_final_cleaned_new <- dplyr::select(meteo_final_cleaned_new, -rownr)
```
62

**Tab. S6:** R-Script used to treat the raw data from the CTD probe.

```
ctd <- read_table2("profiler.txt")
ctd <- dplyr::select(ctd, -Date)
ctd <- dplyr::rename(ctd, c("DATE" = "&", "TIME" = "Time"))
ctd$TIME <- as.character(ctd$TIME)

# order by date and time
ctd <- unite(ctd, timestamp, c(DATE, TIME), remove=F)
ctd$timestamp <- as.POSIXct(as.character(ctd$timestamp), format =
"%d/%m/%Y_%H:%M:%OS")
ctd <- ctd[with(ctd, order(timestamp)),]

ctd$DATE <- as.POSIXct(as.character(ctd$DATE), format = "%d/%m/%Y")
ctd$TIME <- as.POSIXct(as.character(ctd$TIME), format = "%H:%M:%OS")

# add hour
ctd$hour <- hour(ctd$TIME); ctd$hour <- as.numeric(ctd$hour)

ctd <- dplyr::select(ctd, -c(TIME))

# exclude negative depths + one datapoint with all values = 571
ctd <- dplyr::filter(ctd, Press >= 0, Press <= 18)

ctd <- dplyr::rename(ctd, c("date" = "DATE"))
ctd <- unite(ctd, date_hour, c(date, hour), remove=F)

# Sal not relevant in Freshwater + Con20 calculated from CondF
ctd <- dplyr::select(ctd, -c(Sal))

# change unit from uS to mS
ctd$Cond[ctd$Cond > 1] <- ctd$Cond/1000

# exclude negative depths + one datapoint with all values = 571
ctd <- dplyr::filter(ctd, Press >= 0, Press <= 18)

# take only full profiles and filter additional profiles out that are added after a full profile

# order again by date and time
ctd <- ctd[with(ctd, order(timestamp, Press)),]

# make a list of all profiles
ctd_profiles <- ctd %>%
  dplyr::group_by(date, hour) %>%
  dplyr::summarise(n_profile = n(),
                   max_depth = max(Press))
ctd_profiles <- unite(ctd_profiles, date_hour, c(date, hour), remove=F)

ctd_final <- ctd[0,]
for(i in 1:nrow(ctd_profiles)){
  # take each profile
  data <- dplyr::filter(ctd, date_hour == ctd_profiles$date_hour[i])

  data_copy <- data[1,] # save the first measurement
```

```
  if(nrow(data) > 1){
    for(n in 2:nrow(data)){ # loop through all other measurements
      # keep adding data as long as Press is still larger than the max. copied
      if(data$Press[n] > max(data_copy$Press)){
        data_copy <- bind_rows(data_copy, data[n,])
        # if a depth is measured twice, delete the second measurement
      } else if (data$Press[n] == data$Press[n-1]){
        next
        # if the profile starts over, break and delete all further measurements
      } else {
        break
      }

    }
  } else {
    next
  }

  ctd_final <- bind_rows(ctd_final, data_copy)

}

colnames(ctd_final) <- c("Press", "Temp", "CondF", "O2","O2ppm","pH", "PAR", "Chla", "Turb",
"PHY", "timestamp","date_hour", "date","hour")

ctd_final$timestamp <- as.character(ctd_final$timestamp)
ctd_final$date <- as.character(ctd_final$date)

# Press: 0 < x < 19 -> already cleaned
# pH: leave like it is (calibrated regularly)
# Temp: usually fine but generally 0 < x < 40
ctd_final$Temp[ctd_final$Temp < 0] <- NA
ctd_final$Temp[ctd_final$Temp > 50] <- NA
# PAR: 0 < x < n
ctd_final$PAR[ctd_final$PAR < 0] <- 0
# O2
ctd_final$O2ppm[ctd_final$O2ppm < 0] <- NA
ctd_final$O2[ctd_final$O2 < 0] <- NA
ctd_final$O2ppm[is.na(ctd_final$O2)] <- NA # as NA when O2 saturation is NA as ppm is
calculated through saturation
ctd_final$O2[ctd_final$O2 > 300] <- NA
# Chla
ctd_final$Chla[ctd_final$Chla < 0] <- NA
ctd_final$Chla[ctd_final$Chla > 200] <- NA
ctd_final$Chla[ctd_final$Chla == 1000] <- NA
# phy in all versions: 0 < x < n
ctd_final$PHY[ctd_final$PHY < 0] <- NA
ctd_final$PHY[ctd_final$PHY == 1000] <- NA
# Turb: sometimes 1000 as fail -> 0 < x < 999
ctd_final$Turb[ctd_final$Turb < 0] <- NA
ctd_final$Turb[ctd_final$Turb > 200] <- NA
ctd_final$Turb[ctd_final$Turb == 1000] <- NA
# CondF: do nothing to those values
```

```
ctd_final$date <- as.POSIXct(as.character(ctd_final$date), format = "%Y-%m-%d")

ctd_final <- ctd_final %>%
  dplyr::relocate(c(timestamp, date, hour, date_hour), .before = Press)
```

**References**

1. Olli, K. Diel vertical migration of phytoplankton and heterotrophic flagellates in the Gulf of Riga. *J. Mar. Syst.* **23**, 145–163 (1999). doi:10.1016/S0924-7963(99)00055-X.
